# Sharp wave ripples in macaque V1 and V4 are modulated by top-down visual attention

**DOI:** 10.1101/2022.03.14.484243

**Authors:** Jafar Doostmohammadi, Marc Alwin Gieselmann, Jochem van Kempen, Reza Lashgari, Ali Yoonessi, Alexander Thiele

**Affiliations:** Department of Neuroscience and Addiction studies, School of Advanced Technologies in Medicine, Tehran University of Medical Sciences, Tehran, Iran; Biosciences Institute, Newcastle University, NE1 7RU, Newcastle upon Tyne, United Kingdom; School of cognitive sciences, Institute for Research in Fundamental Sciences, IPM, Tehran, Iran; Institute of Medical Science and Technology, Shahid Beheshti University, Tehran, Iran

**Keywords:** Sharp Wave Ripple, Attention, Non-human primate, Visual cortex

## Abstract

Sharp-wave ripples (SWRs) are highly synchronous neuronal activity events. They have been predominantly observed in the hippocampus during offline states such as pause in exploration, slow-wave sleep and quiescent wakefulness. SWRs have been linked to memory consolidation, spatial navigation, and spatial decision-making. Recently, SWRs have been reported during visual search, a form of remote spatial exploration, in macaque hippocampus. However, the association between SWRs and multiple forms of awake conscious and goal-directed behavior is unknown. We report that ripple activity occurs in macaque visual areas V1 and V4 during focused spatial attention. The frequency of ripples is modulated by characteristics of the stimuli, by spatial attention directed toward a receptive field, and by the size of the attentional focus. Critically, the monkey’s reaction times in detecting behaviorally relevant stimulus changes was affected on trials with SWRs. These results show that ripple activity is not limited to hippocampal activity during offline states, rather they occur in the neocortex during active attentive states and vigilance behaviors.

## Introduction

Hippocampal sharp-wave ripples (SWR, ripples) are large amplitude deflections (sharp-waves) of the local field potential (LFP) in the hippocampus of rodents, humans and non-human primates that are associated with a brief fast oscillatory pattern (ripple). Across a variety of species, ripple oscillations vary in the frequency from 140-200 Hz in rodent and 80-180 Hz in non-human primates and humans (1–6). SWRs occur at approximately 0.5 Hz in the hippocampus, 0.1 to 0.5 Hz in posterior parietal, retrosplenial, cingulate and at 0.05 Hz in somatosensory, motor and visual cortices during non–rapid eye movement (NREM) sleep (7). Electrophysiological and neuroimaging studies in rodents and monkeys have indicated that during hippocampal SWRs 15% of hippocampal pyramidal cells discharge synchronously, which triggers activation in cortical areas, but suppression in midbrain and brain stem regions (2, 3).

Ripples support memory consolidation by transferring information acquired during waking to cortical networks during sleep and quiescence (7–9). Consolidation occurs through temporal replay of event related activity in hippocampus during ripples (10–15). SWRs are also predictive of future trajectory and performance on a trial-by-trial basis during spatial navigation tasks (16, 17, 17, 18). Finally, they have been implicated in the correct temporal sequencing of place cell activity preceding novel spatial experiences (preplay) (19).

While most studies based on data from rodents argue that hippocampal SWRs are pronounced during offline states (20, 21), they occur during awake states in humans (22), as well as non-human primates during visual search and goal-directed visual exploration. These SWRs are termed exploratory SWRs (23, 24). SWRs occurrence in the hippocampus of monkeys and humans is increased when the gaze of subjects is focused near a target object during search or when patients observe familiar pictures of scenes or faces (5, 23, 25). For the latter, ripple rates also increased during free recall along with a high-frequency band activation around the time of ripple in visual cortex, suggesting a role of SWRs in activating visual cortex during episodic and semantic memory retrieval (5, 19, 23, 25–28). However, the prevalence and role of SWRs in neocortex is poorly explored. To fill this gap, we recorded LFPs and spiking activity in visual area V1 and V4 of two male macaque monkeys performing a cued spatial attention task. Ripple activity was detected in both regions, ripples occurred more often when monkeys deployed attention to the receptive field of the recorded neurons, and ripple occurrence in V1 was predictive of better behavioral performance. These data suggest that ripples occur in cortical sensory areas and are involved in cognitive functions beyond memory consolidation and retrieval.

## Results

Monkeys performed a covert spatial attention paradigm (Methods and figure 1). We recorded 26 and 17 sessions in monkeys 1 and 2 respectively with overlapping receptive fields between V1 and V4 (Table S1, supplementary materials and methods). We analyzed three main epochs for SWR occurrence, namely (i) pre-cue: a 200 ms period after fixation before cue onset, (ii) post-cue (800 ms period after cue offset until stimulus onset) and (iii) the sustained response period corresponding to 300-750 ms after stimulus onset (Supplementary figure S1B).

**Figure 1.**
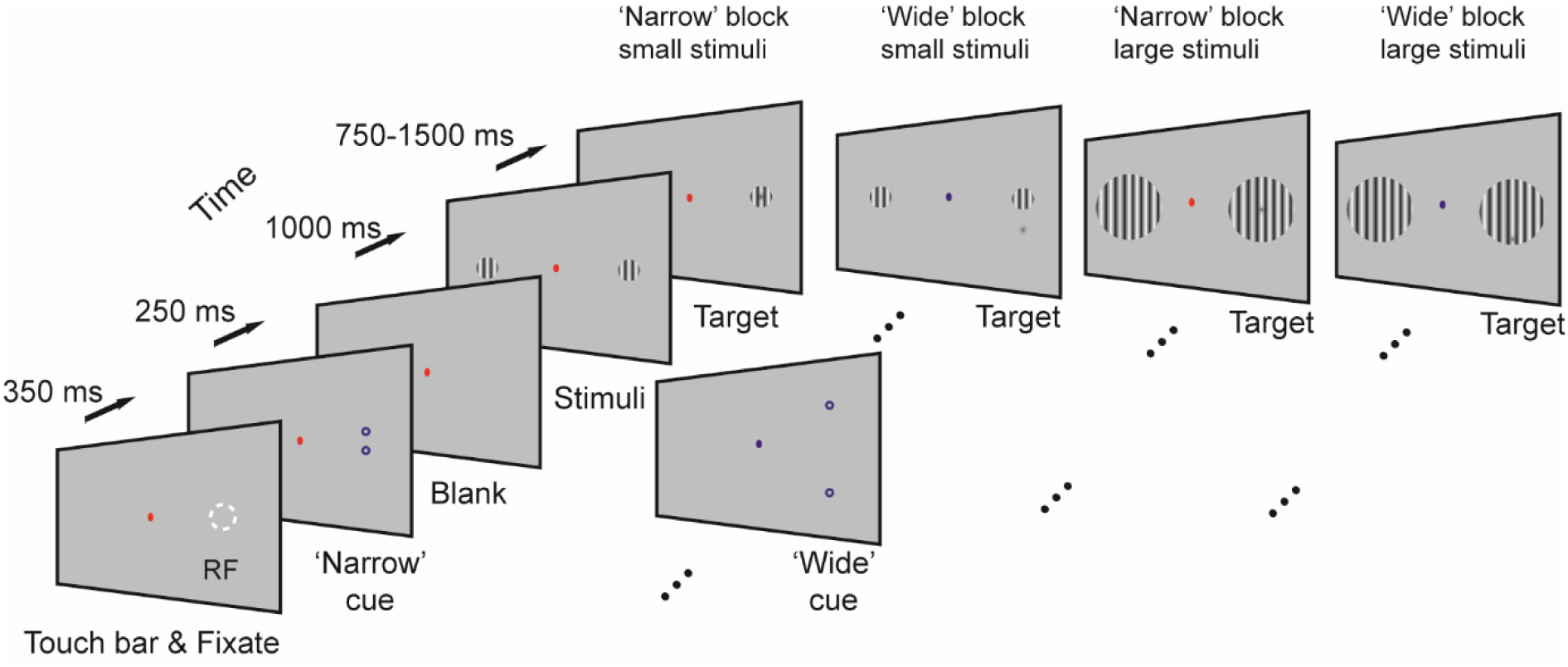
Behavioral paradigm. Monkeys were trained to touch a touch bar, which triggered a fixation point to appear on the screen. Following fixation onset, two cues (small annuli) were presented for 250 ms which indicated which side of the screen was behaviorally relevant. The distance of the two rings, as well as the color of the fixation spot, indicated whether the trial belonged to a narrow or wide focus of attention condition. 1000 ms after cue offset, two static grating were presented, one centered on the receptive field (white dotted circle, for illustrative purpose, not present on the screen during the experiment), the other grating appeared equidistant to the fixation spot in the opposite visual hemifield. After 750-1500 ms a dimming spot appeared in the cued location, upon which the monkey had to release the touch bar with 500 ms to receive the fluid reward. In narrow blocks the dimming spot was generally presented centered on the target stimulus while in wide blocks it was generally presented at an unpredictable location offset from the stimulus center.

### Ripple attributes across task epochs

The total number of ripples detected in V1 were 166, 688, 7768 for pre-cue, post-cue and sustained periods, respectively. In V4 the relevant numbers were 642, 2740, and 9545 for pre-cue, post-cue and sustained periods. Example trials with ripples observed during the task epochs are shown in figure 2A-E. Spectrograms of ripples revealed a high-frequency component oscillation (100-180 Hz) in V1 as well as in V4 (figure 2E-G and supplementary figures S2&S3). Ripples in V1 showed lower peak frequency during the sustained period compared to pre and post cue periods, while in V4 ripples had higher peak frequency during the sustained period (supplementary figure S3, V1; χ _(2)_^2^ = 15.9365, p= 0.0003, V4; χ _(2)_^2^ = 145.7693, p = 2.2213e-32, Kruskal-Wallis test). The majority of ripples had durations between 30-60 ms. On average, the mean and SEM of durations of ripples were 45.19 ± 1 ms, 43.95 ± 0.5 ms and 47.67 ± 0.5 ms in V1 for pre-cue, post-cue and sustained periods respectively. In V4, these durations were 47.6 ± 1 ms, 45.2 ± 0.5 ms and 47.8 ± 1 ms (supplementary figure S4). Ripples during the sustained period were longer than in other periods in both areas (V1; χ _(2)_^2^ = 26.8762, p= 1.4585e-06, V4; χ _(2)_^2^ = 59.7597, p = 1.0552e-13, Kruskal Wallis test).

**Figure 2.**
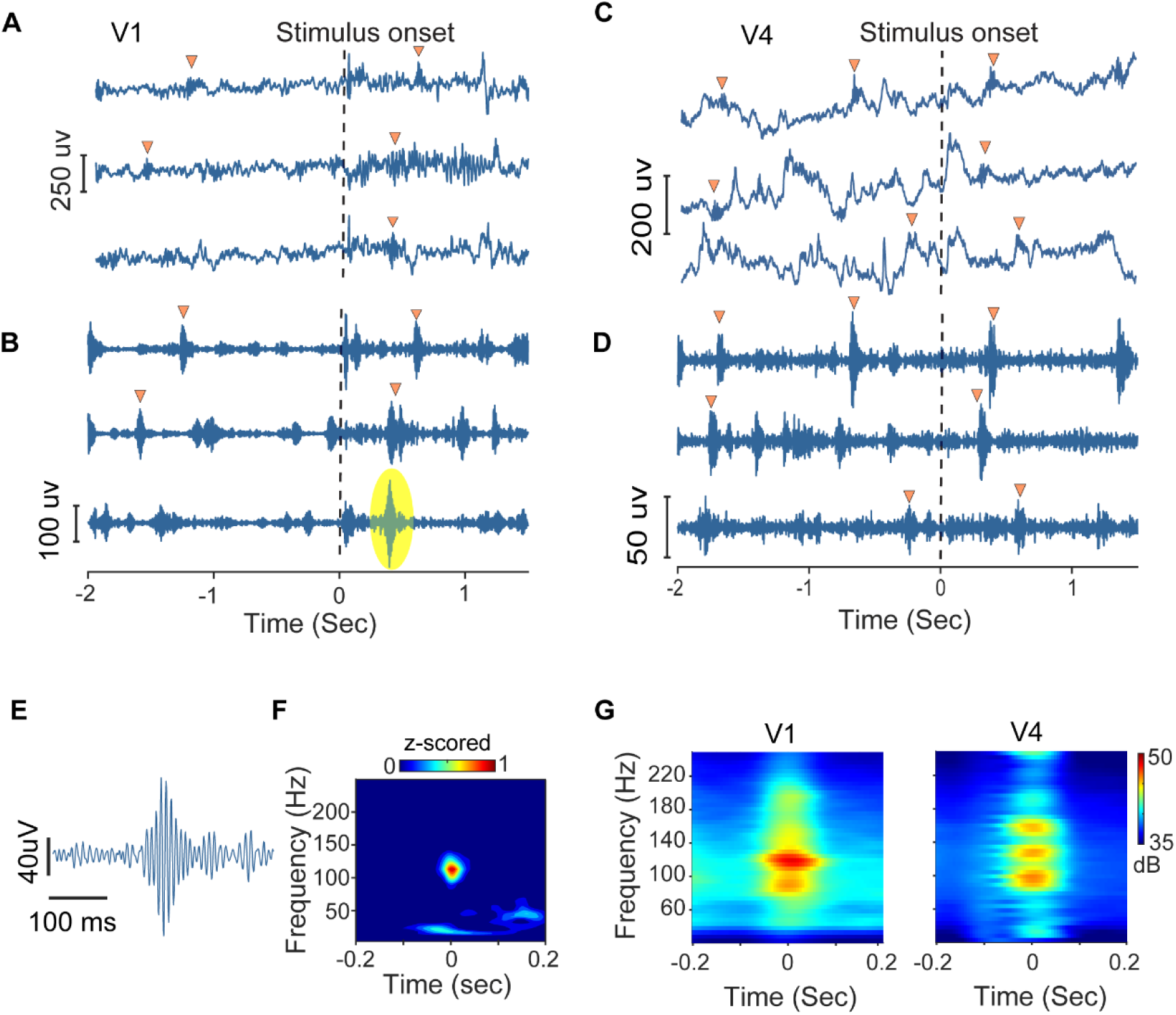
Example ripples recorded during task epochs time locked to stimulus onset. A, C) Bipolar re-referenced LFPs demonstrating occurrence of ripple in the task epochs during visual attention in V1 and V4. B, D) respective ripple band voltage traces (bandpass filtered 80-250 Hz). Time zero and dotted line indicate stimulus onset. Ripples are shown with yellow triangles. E, F) Representative magnified ripple trace and spectrogram indicated with a yellow oval in B. Color bar represents z-scored power. G) Overall average of Hanning windowed taper spectrogram showing time-frequency distribution of ripples identified in V1 and V4. Color maps reflect power of ripples in decibel (dB).

### Ripples and cortical layers

We next analyzed whether ripple rates differed between cortical layers of visual cortex. The depth of recording contacts was determined by multi-unit response latency and current source density analysis (CSD) of LFPs as described in methods. Supplementary figure S5 illustrates the CSD profile and laminar alignment to characterize the layers (methods). We assigned layers I-III to supragranular, layer IV to granular, and layers V and VI to infragranular layers. As the thickness of layers varies within and between visual areas, we divided ripple rate over the number of the contacts embedded in the corresponding layer. Ripple rates were similar in supra-granular, granular and infra-granular layers of V4 (F (2,126) = 0.57, p = 0.56, one-way ANOVA). In V1, the granular layer showed higher ripple rates than the supragranular and infragranular layers (one-way ANOVA over pooled data and layer as factors, F (2,119) = 9.44, p = 0.0002, post-hoc Tukey: supragranular vs. granular, mean difference = -0.04, 95% C.I = [-0.06 -0.01], p=0.002, granular vs. infragranular: mean difference = 0.04 C.I = [0.01 0.07], p = 0.003), while the ripple rate did not differ between supra-granular and infra-granular layer (supplementary figure S6)

### Effect of stimulus size on ripple events

To quantify ripple rates as a function of stimulus size, ripple events were compared when large and small stimuli were presented. Small stimuli were associated with higher ripple rates than large stimuli. This effect was observed for both areas and it was consistent across subjects. Mean and SEM of ripple rates in V1 and V4 for small stimuli were 0.09 ± 0.006 Hz and 0.1 ± 0.007 Hz, while they were 0.04 ± 0.002 and 0.04 ± 0.003 Hz for large stimuli (V1: Z = -4.7, p = 1.5982e-06; V4: Z = -4.4, p = 7.9053e-06, Wilcoxon signed rank test, pooled for both monkeys, figure 3A). The effects of stimulus size on ripple rate were not an artefact of centering the stimuli on the (smaller) V1 RFs, potentially causing offsets between V1 and V4 RF stimulation, and possibly not stimulating some V4 RFs when small stimuli were used. Firstly, if that was the case we would have found lower ripples rates comparable to pre-stimulus periods for small stimuli in V4, not higher ripple rates. Secondly, in monkey 1, most recordings had V1 RFs that were completely covered by all V4 RFs, i.e. the small stimuli covered both RF locations well (supplementary figure S7). For these recordings (n=18) we found the same effect of stimulus size on ripple rates as for the entire set of recordings (supplementary figure S8). For additional details see supplementary information (SI Ripple rate and RF overlap).

**Figure 3.**
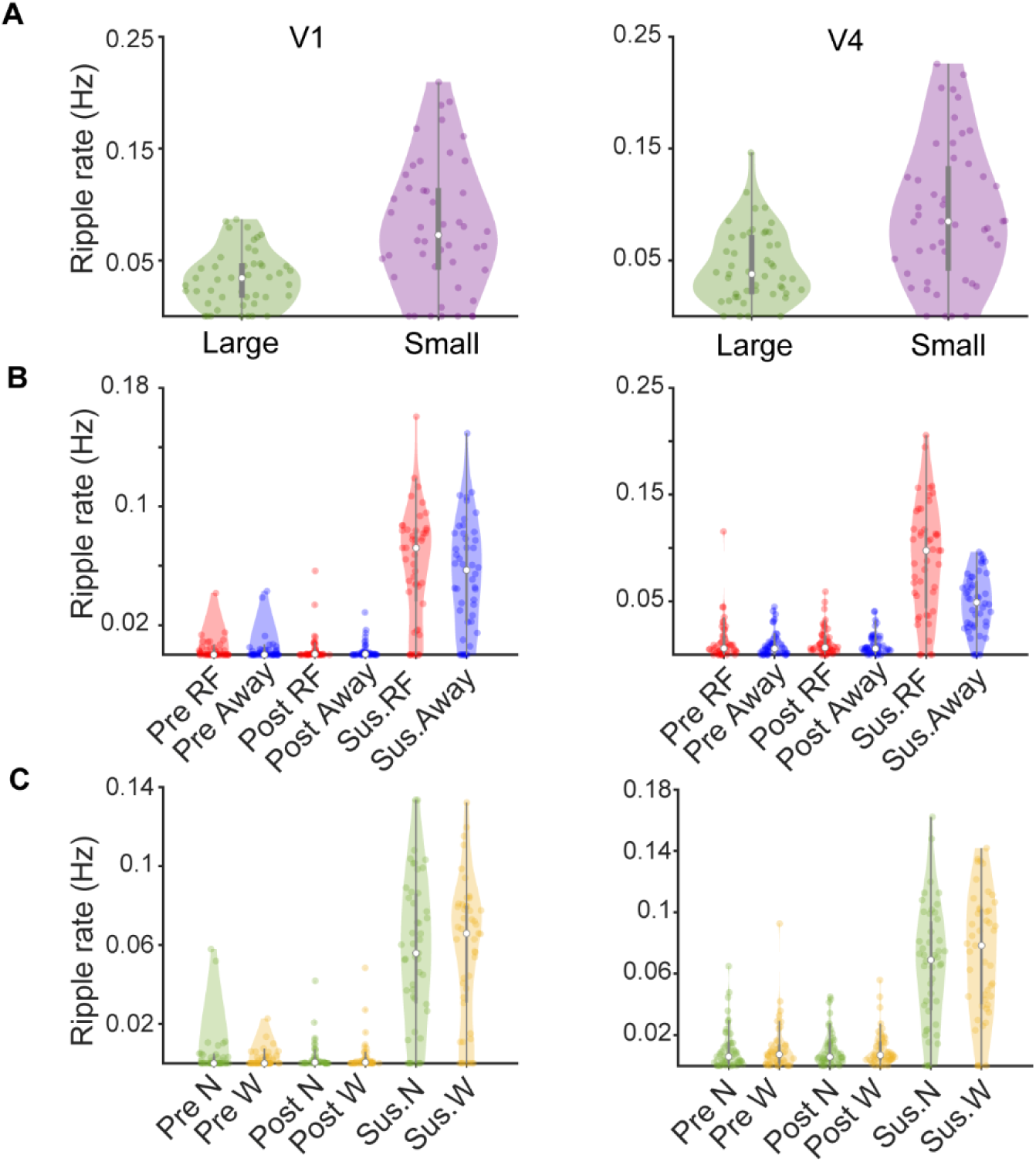
Ripple rate as a function of stimulus size, attention to receptive field and size of focus of attention. A). Effect of stimulus size on ripple rate. Left column shows ripple rate in V1 for small and large stimuli. Right column shows ripple rate as a function of stimulus size for V4. B) Comparison of ripple rate for different attention conditions (attend RF vs. away) for 3 different trial periods (pre-cue, post-cue, post stimulus sustained period). Left column shows V1 data (red- attend RF, blue-attend away). Right column shows data for V4. C). Ripple rate as a function of size of attentional focus (narrow vs. wide) for the three trial periods (pre-cue, post-cue, post stimulus sustained period). Ripple rate did not differ between attentional focus size in V1 (left) for either monkey. In V4 wide focus of attention triggered higher ripple rates than narrow focus of attention. N and W indicate narrow and wide blocks of attention.

### Relation of ripples to attention states, and possible interactions with stimulus properties

We next explored whether ripple rate changed with task conditions. We detected ripples in task epochs as well as inter-trial intervals. The mean and SEM of ripple rate across all trials in V1 was 0.003 ± 0.009 Hz for pre- and post-cue and 0.056 ± 0.002 Hz during sustained periods. In V4 the mean and SEM of ripple rate was 0.01 ± 0.001 Hz during pre- and post-cue and 0.068 ± 0.003 Hz during the sustained period. Ripple rate in inter-trial intervals was 0.02 ± 0.002 Hz for both regions.

To examine whether attention influenced ripple rate, we computed ripple rate in trials when monkey’s attention was directed toward vs away from the receptive field (referred to as attend RF vs. attend away). Ripple rate did not differ between attention to RF and away during the pre- and post-cue periods in V1 (pre-cue: Z = 1.06, p = 0.28, post-cue: Z = 0.36, p = 0.71, Wilcoxon test) or V4 (pre-cue: Z = 0.95, p = 0.33, post-cue: Z = 1.36, p = 0.08, Wilcoxon test). However, results differed for the sustained period. Here, the mean and SEM of ripple rate was higher for attend RF conditions in V1 (0.06 Hz in attend RF vs. 0.05 Hz in attend away, Wilcoxon sign rank test; V1: Z = 2.4732, p = 0.0134, figure 3B). In V4 ripple rate was increased by spatial attention during the sustained period (0.09 for attend RF vs 0.04 for attend away, Wilcoxon sign rank test; Z = 5.66, p <0.001, figure 3B). Figure 3 shows the overall distribution of ripple rates for the effects of stimulus size, locus of attention, and focus of attention across sessions.

While the above analysis shows that attention affected ripple rates during the sustained period, it did not dissect effects according to different stimulus and attentional focus conditions. To assess whether effects of attention depended on stimulus size and/or the size of the attentional focus, we calculated ripple rates for all triplet-wise conditions (eight possible permutations: attend-RF/small-stimuli/narrow-focus, attend-RF/small-stimuli/wide-focus, attend-RF/large=stimuli/narrow-focus, ….) for each electrode for each session, and then averaged ripple rates across electrodes for a given session. This yielded eight values corresponding to the eight possible condition combinations for each session. We then used a 3-factor repeated measures ANOVA, to determine main effects of stimuli, attentional location and attentional focus, and possible interactions on ripple rates in V1 and V4.

#### Effects in area V1

are shown in figure 4A-F. The 3-way ANOVA revealed that stimulus size (p<0.001, figure 4A) and attention (p=0.014, figure 4B) had a main effect on ripple frequency in V1 data, while the focus of attention did not have a main effect (p=0.902, figure 4C). Details about the statistics (df, F-values, …) are listed in supplementary table S2. Small stimuli resulted in higher ripple rates than large stimuli, and attention to the RF resulted in higher ripple rates. Additionally, we found a significant stimulus*attention (location) interaction (p=0.035), a significant stimulus*attentional focus interaction (p<0.001), and a significant attentional location*focus interaction (p=0.009). The effect of attentional location on ripple rate was significant when small stimuli were presented (p=0.002, post-hoc sign rank test, figure 4D), but not when large stimuli were presented (p=0.293, post-hoc sign rank test, supplementary figure 4D). For the stimulus*attentional focus interaction we found that wide foci of attention yielded higher ripple rates than narrow foci when small stimuli were presented (p=0.005, post-hoc sign rank test, figure 4E). The opposite was the case when large stimuli were presented (p<0.001, post-hoc sign rank test, figure 4E). This difference itself was significant (p<0.001, sign rank test based on pairwise differences for the two groups, figure 4E). Finally, the locus of attention* focus of attention interaction was a result of slightly increased ripple rates for wide focus of attention under attend RF conditions, and slightly decreased ripple rates for wide focus of attention conditions under attend away conditions (always compared to narrow/small focus). However, neither of these two comparisons were significant by themselves (post-hoc sign rank test, details in figure 4F).

**Figure 4.**
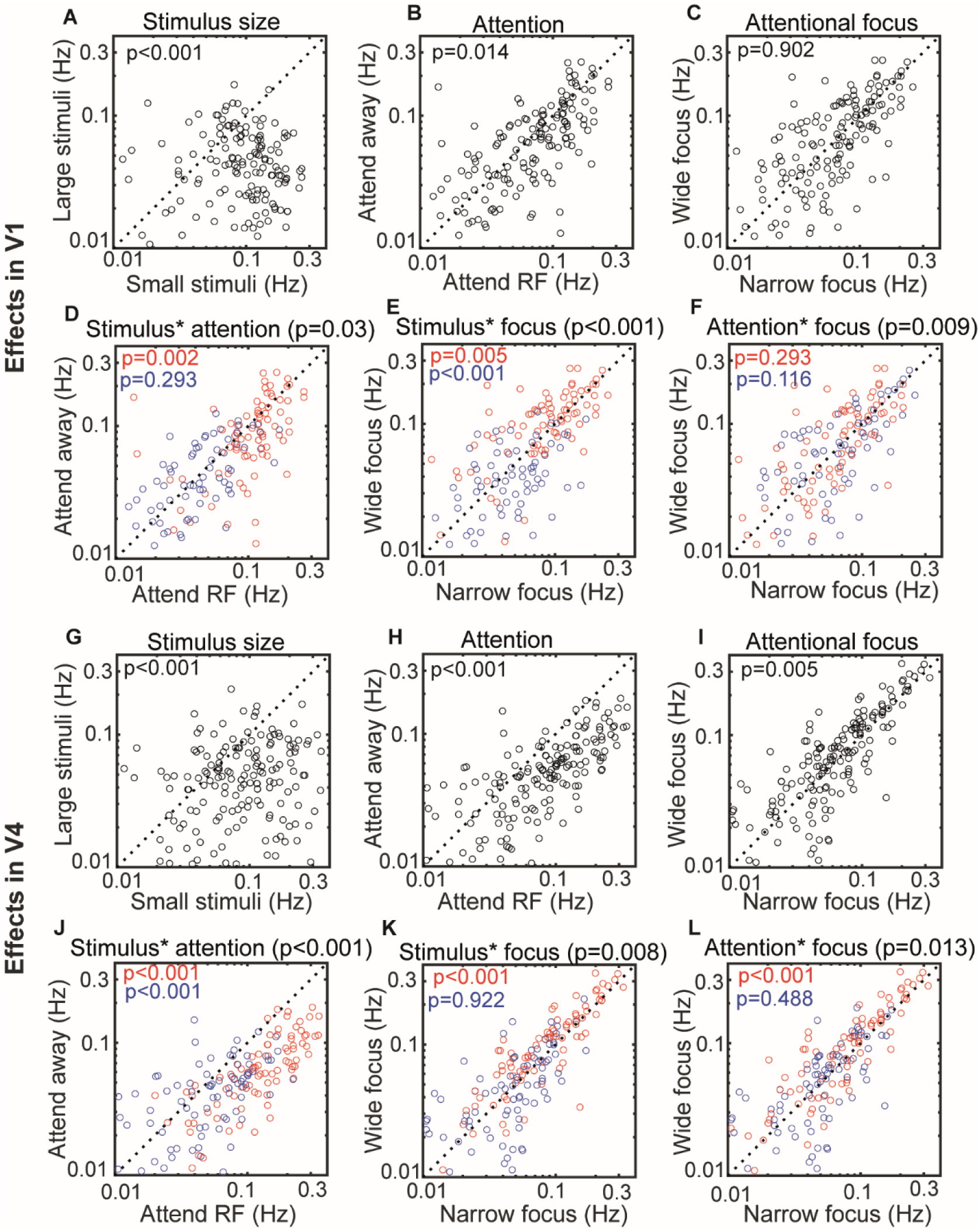
Ripple rate in areas V1 and V4 as a function of stimulus size, location of attention, and attentional focus, as well as possible pairwise interactions. A) Main effect of stimulus size on ripple rate. B) Main effect of attention (location) on ripple rate. C) Main effect of focus of attention on ripple rate. D) Interaction between stimulus size and attention location on ripple rate. Red: ripple rate for small stimuli for attend RF vs attend away conditions. Blue: ripple rate for large stimuli for attend RF vs attend away conditions. E) Interaction between stimulus size and focus of attention on ripple rate. Red: ripple rate for small stimuli for narrow focus vs wide focus conditions. Blue: ripple rate for large stimuli for small focus vs wide focus conditions. F) Interaction between attentional locus and focus of attention on ripple rate. Red: ripple rate for attend RF conditions for narrow focus vs wide attentional focus. G-L) Same as A-F but for area V4. Blue: ripple rate for attend away conditions for narrow focus vs wide attentional focus. Numbers indicate mean ± SEM and p-values respectively. Color coded p-values are from post-hoc sign rank tests.

#### Effects in area V4

The effects of stimulus size, attention location and attentional focus and possible interactions for area V4 are shown in figure 4G-L. The 3-way ANOVA revealed that stimulus size (p<0.001, figure 4G), attention (p<0.001, figure 4H), and attentional focus (p=0.005, figure 4I) had a main effect on ripple frequency in V4 (statistical details are listed in supplementary table S3). Small stimuli resulted in higher ripple rates than large stimuli. Attention to the RF (compared to attend away) resulted in higher ripple rates, and a wide focus of attention (compared to narrow focus) resulted in higher ripple rates. Additionally, we found a significant stimulus*attentional location interaction (p<0.001), a significant stimulus*attentional focus interaction (p<0.001), and a significant attentional location*attentional focus interaction (p=0.005). Attention to the RF resulted in higher ripple rates than attention away, when small stimuli were presented (p<0.001, sign rank test, figure 4J). While this was also the case when large stimuli were presented (p<0.001, sign rank test, figure 4J), the effect of attention on ripple rates was significantly stronger when small stimuli were presented than when large stimuli were presented (sign rank test on pairwise differences between attend RF minus attend away conditions, grouping variable: small vs. large stimuli, p<0.001, figure 4J). A wide focus of attention resulted in higher ripple rates than a narrow focus of attention, when small stimuli were presented (post-hoc sign rank test, p<0.001, figure 4K), but not when large stimuli were presented (post-hoc sign rank test, p=0.922, figure 4K). This difference itself was significant (p<0.001, sign rank test on differences between narrow and wide focus of attention for small vs. large stimuli). Finally, when attention was directed to the RF a wide focus resulted in higher ripple rates than a narrow focus (p<0.001, post-hoc sign rank test, figure 4L), while no difference for the focus of attention was found when attention was directed away from the RF (p=0.448, post-hoc sign rank test, figure 4L).The difference of attend RF vs. away on attentional focus difference was significant (p=0.009, sign rank test, figure 4L).

### Spiking activity at ripple time

We next compared firing rates during the sustained period at ripple time versus firing rates during trials without ripple for identical conditions (figure 5, supplementary tables S4 and S5 give a breakdown of firing rates for the multiple conditions tested in V1). In V1, the mean and SEM of thresholded MUA_E_ (methods) firing rate during the sustained period without ripples and attend RF trials was 144.6 ± 1.2 Hz, which increased to 164.3 ± 1.5 Hz in trials with ripples. In attend away trials ripples enhanced average firing rates to 165.2 ± 1.3 Hz, while in the same condition without ripples the rate was 140.6 ± 1.4 Hz. Similarly, in V4, the mean and SEM of firing rate during the sustained period in trials without ripples and attend RF was 101.6 ± 0.8 Hz, which increased to 123.4 ± 0.99 Hz with ripples. Ripples in attend away trials resulted in mean firing rates of 109.6 ± 1.2 Hz, while without ripples rates were 85.17 ± 1.02 Hz. Figure 5 gives an overview of firing rates in trials with and without ripple for condition and trial matched data for V1 and V4. To further explore which factors of the task contribute to increased neural activity, we conducted a linear mixed effect model analysis to predict the firing rate based on attention (RF/away), ripple occurrence (presence/absence), focus of attention (narrow/wide) and size of stimulus (large/small). The mixed effects model in V1 revealed that the firing rate was modulated by ripple, focus of attention and stimulus size (ANOVA following mixed effect model: Ripple: F (1, 16578) = 243.1, p < 0.001, Focus: F (1, 16578) = 9.92, p = 0.001, size: F (1, 16578) = 694.97, p < 0.001). Similarly, in V4 firing rate was modulated by ripple, attention to RF and size of the stimulus. (ANOVA following model: Ripple: F (1,18932) = 442.2, p < 0.001, Attention: F (1,18932) = 128.6, p < 0.001, size: F (1,18932) = 160.6, p < 0.001). In addition, firing rate in V4 also depended on an interaction between ripple and attention to RF (F (1, 18932) = 17.04, p < 0.001). Supplementary tables S6 and S7, summarize the results of the linear mixed effect model in V4.

**Figure 5.**
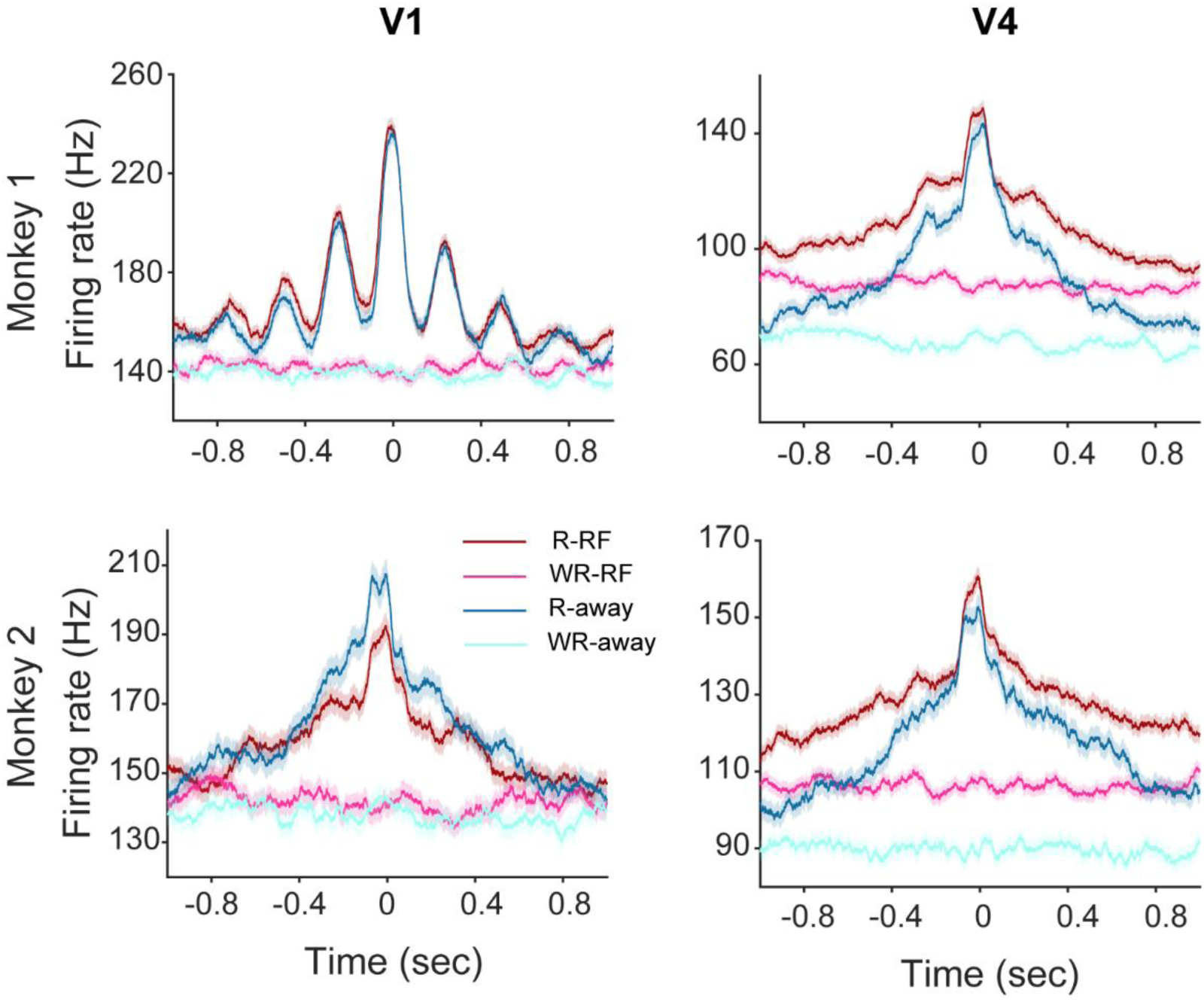
Multi-unit spiking activity with and without ripple events. Mean and SEM of multi-unit spiking activity centered at ripple time in V1 (left) and V4 (right) on trials with and without ripple. Dark red and blue lines show trials with ripple on attend RF/away conditions, pink and cyan are trials without ripples, chosen from the same number of trials as with ripples, under identical conditions (i.e. attend RF/away, narrow wide focus, …), and matched time points within a trial. R-RF, and R-away: ripple during attention to RF and away conditions. WR-RF and WR-away: without ripples during attention to RF and away conditions.

### Cross-correlation of V1-V4 ripples

To assess the temporal association between ripples in V1 and V4, we computed cross correlations between V1 and V4 ripples between all channel combinations, pooled across session (figure 6). We corrected the cross-correlation by subtracting the shuffle predictor from the raw cross-correlation of ripples on each session. If a ripple occurred on any contact in V1 or V4, then the probability of a ripple occurring on that trial in the other area was 0.097. We used area under the cross correlogram (AUC) relative to time zero as indications of lead-lag relationships. The AUC values did not differ for lead or lag periods, i.e. there was no evidence for a lead-lag relationship between V1 and V4 ripples (Monkey 1: Z = -1.3857, p = 0.1658, monkey 2: Z = -2.5860, p = 0.7354, Wilcoxon signed rank test).

**Figure 6.**
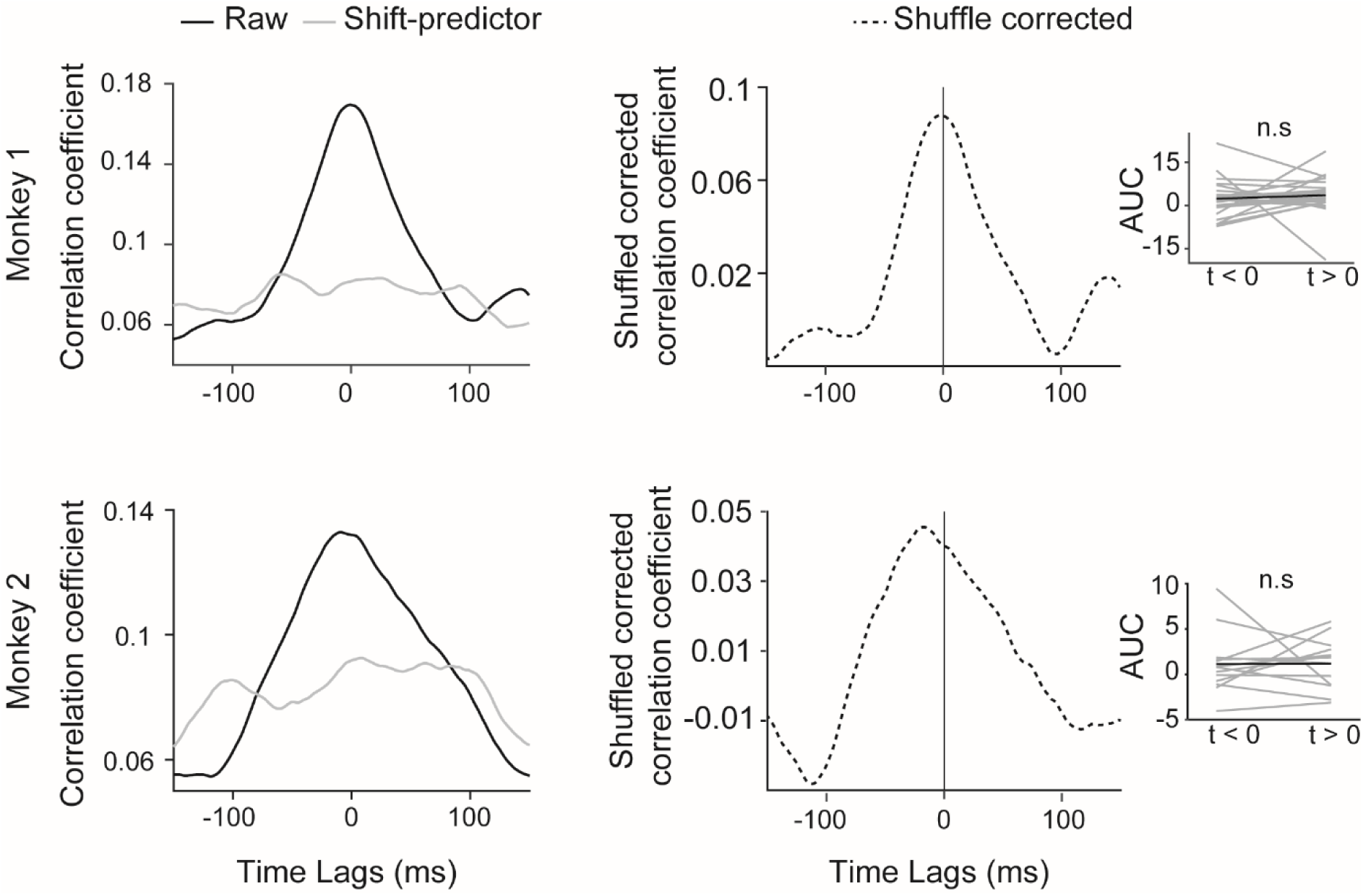
Temporal cross-correlation between ripples in V1 and V4. Black curves in the left panel represent temporal cross-correlation between V1 and V4 ripples. Gray curves correspond to cross-correlation of shuffled trials used for correcting the raw cross-correlation. Dotted curves in the right panel show shuffle corrected correlation of V1 and V4 ripples. Insets indicate area under the curve for lead (AUC to left of ripple center) and lag AUCs.

### Cross-frequency power coupling between ripples

Next, we analyzed whether ripple activity was coordinated between layers, between areas, and whether ripple activity showed specific cross frequency coupling to other spectral frequency bands. To examine this, we estimated the time-frequency spectrogram of V1 and V4 LFPs using either V1 or V4 ripples as trigger events. Figure 7 depicts the average of power coupling across all channel combinations of V1 and V4 using V1 ripples as the trigger. Power-power coupling was dominant for the ripple and gamma band between V1 and V4 (at α = 0.05, p = 0.0065, FDR correction). However, power correlation was not significant between areas when V4 ripples served as trigger. This shows that ripples in V1 affect LFP spectral power in V4, i.e. in the feed-forward, but not the feedback direction.

**Figure 7.**
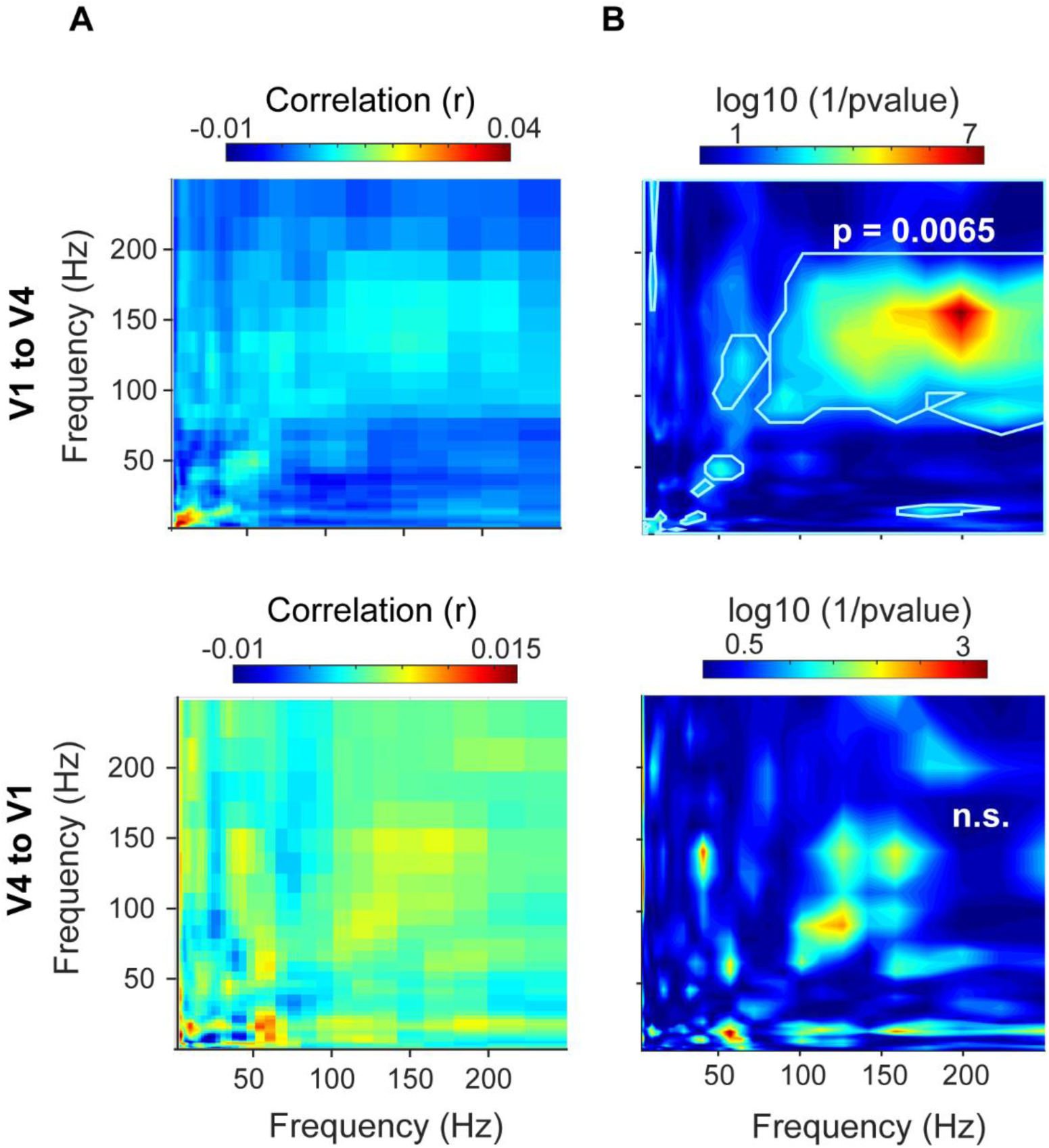
Modulation of spectral power between V1 and V4 around the time of ripple events. A) Upper panel shows average comodulogram of V1 and V4 when V1’s ripples served as trigger. Comodulogram demonstrated frequency coupling between V1 and V4 at ripple and gamma frequency band. Lower panel represents average corrected comodulogram of frequency coupling between V4 and V1 where ripples in V4 served as trigger. B) Logarithm with base 10 of 1/p-values after FDR correction for V1 (upper) and V4 (lower). Points inside solid light line represent frequency bands that showed significant coupling at V1’s ripple episodes.

To determine layer-specific coordination, we analyzed the power correlation between pairs of contacts in the three laminar compartments within each area, using ripple events as triggers as described above. The comodulogram profile for V1 showed that the power-power coupling between supragranular to supragranular, granular to supragranular and infragranular to supragranular was higher in the ripple band. This effect was pronounced in the reverse direction as well (supplementary figure 9). In V4 the power coupling was significant when infragranular served as a trigger for the supragranular as well as supragranular to supragranular contacts, with no other interactions (supplementary figure 9).

### Ripple rate and reaction time

Finally, we asked whether ripple occurrence during attention was linked to the RT. We computed RTs in different task conditions across all trials of both subjects. The RTs were classified by no ripple, ripple occurrence in V1 trials, ripple occurrence in V4 trials, and finally ripples in both areas. Regardless of task conditions, the mean and SEM of z-scored RTs on trials without ripples was 0.003 ± 0.01, whereas on trials when ripples only occurred in V1, the mean and SEM was -0.02 ± 0.01. The RT in trials where ripples only occurred in V4 was 0.02 ± 0.02. When ripples occurred in V1 and V4, mean and SEM of RT was -0.02 ± 0.02. These results suggested that ripple occurrence in V1 and both areas decreased RTs while ripple occurrence only in V4 increased RTs of the subjects (supplementary figure S10). To probe this further, a linear mixed effect model was used to identify which task variables were significant predictors of RT. Attention to/away from RF, focus of attention (wide-narrow), stimulus size and ripple occurrence (no ripple, V1 or V4 ripple, ripple on both areas), were used as predictors. The result of the model showed an interaction of trials without ripple, narrow focus of attention and large stimuli. Specifically, these conditions increased the reaction time of the monkeys (F (3, 15458) = 2.2, p = 0.02). This result suggests ripple occurrence, especially on V1 or both areas contribute to modulate the RT of the subjects not in independent manner but with other task variables. Refer to supplementary table of 8 and 9 for details of the linear model on RTs.

## Discussion

Here we provide insight into ripple activity in the visual cortex during spatial attention. We demonstrated that ripples occurred during different epochs of top-down attention in visual areas V1 and V4 of macaque monkeys, with increased ripple rates when monkey’s attention was directed to the RF in both regions. Our results highlight four new aspects of SWR functions and their association to visual attention. First, ripple rate was higher when small stimuli rather than large stimuli were presented. Second, top-down attention enhanced the ripple rate in V1 and V4. Third, there was a power correlation in the ripple band within and between striate and extrastriate area V4. Finally, the presence of ripples during the sustained periods modulated the reaction time of the monkeys.

SWRs occur most frequently during quiescence and non-REM sleep in the hippocampus of rodents where they are important for memory consolidation (10, 12, 29) and less frequent during waking periods where they appear to be important for memory-guided decision making (30–32). Additionally, SWRs occurred at specific points in maze navigation where they predict future navigational behavior (17, 33–35). When rodents learn a navigation task, SWR rate increased before successful trials during the learning period (18). SWRs have recently been reported in the primate hippocampus, where they occur during active visual search and memory retrieval (23, 25) suggesting they are required to integrate immediate past experiences with current rules (36). Most previous studies have reported SWRs in the hippocampus, but they have recently also been reported to occur in rodent association cortex (7) during non-REM sleep and quiet wakefulness. In humans increased ripple rates occur in hippocampus during autobiographical episodic and semantic memory recollection tasks (5, 27). To the best of our knowledge, our study is the first to show modulation of ripple rates by spatial attention, and by visual stimulus characteristics in the striate and extrastriate areas in the non-human primate, during a cued covert spatial attention task. Critically, attention directed to the receptive fields of the neurons increased ripple rates and, ripple occurrence in V1 or V1 and V4 during a trial resulted in faster reaction times, while the opposite was the case when ripples occurred only in V4. These effects depended on the attentional focus size, whereby ripples decreased RTs when the attentional focus was wider. Many studies implicate SWRs in memory retrieval and planning. A possible interpretation of SWRs in visual cortex during spatial attention is that the subjects had to memorize the location of a spatial cue to detect the target to release a touch bar. Thus, ripples in our study might indicate a form of memory recollection to accomplish the task. However, memory in our task would probably be working memory, rather than forms of spatial (or even episodic) memory as often probed in previous studies. Studies in rodents uncovered that ripples are prevalent at reward sites of a navigation task (37). Since in our paradigm, attention was linked to likelihood of receiving a reward, this might be an explanation of the higher ripple rate during the sustained period with attend RF conditions. A higher ripple rate in wide blocks particularly in V4 is also consistent with these arguments, because on wide blocks the uncertainty of the subjects was higher, therefore they may demand more attention to receive a reward, and wide blocks elicited higher ripple rates. The latter presumable also has a covert search component (across the wide focus of attention), and it might thus be linked to increased ripple rates seen in primate hippocampus during active visual search (23, 25, 38). However, it appears that ripples in V1 and V4 may serve different functions and may engage different networks. Ripples in V1 during attend RF trials reduced RTs, while ripples in only V4 during attend RF trials increased RTs. There is a wealth of research indicating high frequency oscillation are involved in the feedforward and intra-cortical communication (39–42), Ripples in V1 could serve this function, possibly maintaining specific communication subspaces (43) required for efficient task performance. The fact that cortical ripple band activity in V1 coupled with gamma and ripple band activity in V4 might be a signature of this. The reason why ripples in V4 increased RTs, suggesting reduced task performance, is difficult to reconcile with this framework. If it were a signature of intermittent cue/task recollection, then this might interfere with efficient task performance (44) and result in slower reactions. Future work is required to explore these possibilities.

## Materials and Methods

### Animals and procedures

Details regarding animals, housing, surgical procedures, animal and ethical licensing, visual stimulation, eye monitoring, recording hardware, and procedures are given in supplementary materials.

### Visuospatial attention task

The monkeys were trained to detect a small, low contrast luminance stimulus appearing on a square wave grating in the peripheral visual field while centrally fixating (figure 1). If the small stimulus occurred at a cued location it is referred to as ‘target’, if it occurred at an uncued location it is referred to as ‘distractor’. Hence the paradigm involved cued spatial attention, whereby the cue indicated also whether the attentional focus should best be narrow or wide (see below for details). Monkeys initiated a trial by holding a touch bar. Subsequently a fixation point (FP, red annulus for attend RF, blue for attend away conditions, 0.17 deg diameter) appeared at the center of the screen on a grey background. A drift in eye position of more than 1.5 or 0.9 deg (monkey 2 or monkey 1) throughout the trial was registered as a fixation error and the trial was terminated. Following a period of initial fixation (monkey 2: 500 ms, monkey 1: 350 ms), a spatial cue (a pair of blue annuli, 0.17 deg diameter) was presented for 250 ms. The spatial cues served two purposes: (1) they indicated in which visual field the target stimulus would later appear, and hence which visual location should be attended to (contralateral vs. ipsilateral to the recording locations, i.e. attend to the RF of recorded neurons vs away), and (2) whether the best strategy would be to employ a narrow or wide attentional focus. If the 2 cue dots were placed close together [cue center distance: 0.2°] it indicated that the target was likely to appear centered on the stimulus (see below), while 2 cue dots placed further apart [cue center distance: 5.0°] indicated that the target likely appeared at 1 of 12 possible positions peripheral to the stimulus center. One second after cue offset, two static circular square wave grating stimuli of either 1° or 5° diameter (duty cycle of 1.5 cycle/degree, contrast 30%, monkey 2, or 40%, monkey 1) were presented. The grating orientation was chosen from one of 6 possible orientations (60° orientation spacing) to match the aggregate preference of neurons at the recording site (the orientation was fixed for a given recording session). One of the gratings was presented on the aggregate visual receptive field, with a focus on centering it on V1 RFs, as these were smaller. The other grating was presented in the opposite visual hemifield, mirrored at the FP. After a randomized period of 0.75-1.0 s or 0.75-1.25 s (monkey 2 and monkey 1 respectively) either the target appeared at the cued stimulus location, or the distractor appeared at the stimulus location in the opposite visual hemifield. The monkey had 500 ms to report (touch bar release) target appearance at the cued location to receive a reward (the amount of reward increased with decreasing reaction time) and ignore distractor appearance. If the distractor appeared at the uncued location, a target appeared at the cued location 1-1.4 s (monkey 2) or 1.5-1.75s (monkey 1) after the distractor had appeared. The target/distractor was a circular patch of Gaussian modulated (σ=0.1°) luminance of variable contrast relative to the parts surrounding it (which could be either the background or grating stimulus, dependent on target location and grating stimulus size). The position of the target (central/peripheral to grating center) was varied block wise. The size of the grating (large/small) was varied block wise in monkey 2, and trial wise in monkey 1. The order of target appearance (in RF first/second) was varied pseudo-randomly trial wise. Narrow and wide focus of attention blocks were additionally coded by the color of the fixation spot (narrow = red, wide = blue). For wide focus of attention conditions, the possible target position was one of at least 12 possible target positions peripheral to the grating center. In a “narrow” block the target appeared in the center of the grating in 80% of the trials. In a “wide” block the target appeared peripheral to the center of the grating in 80% of the trials. In invalidly cued catch trials (7% in monkey 2, 10% in monkey 1) we presented a peripheral target in “narrow” blocks and a center target in “wide” blocks. In blank trials (13% in monkey 2, 10% in monkey 1) no target was shown, and the monkey received a reward for holding on to the touch bar. Reaction times (RT) of monkeys were calculated by subtracting the time of touch bar release from the time of appearance of the target. Due to non-normal distribution of RTs, data were natural log transformed. They were z-scored for each recording session to remove the effect of differential stimulus eccentricity. Details about receptive field mapping are given in supplementary materials.

### Data acquisition

Neurophysiological recordings were performed with laminar silicon electrodes comprising of 16 contacts with a 150 µm spacing between adjacent channels (Atlas Neuroengineering, Belgium). The impedance of the contacts was measured before each penetration and were in the range of 0.5-1.0 MΩ. The probes were mounted on a hydraulic Microdrive (Narishige MO-97A, Japan). The probes were inserted perpendicular to the cortical surface with the intent to yield coverage of all laminae along the gray matter of the cortex.

Extracellular voltage fluctuation of V1 and V4 were acquired simultaneously using a 32-channel digital Lynx acquisition hardware (Neuralynx, USA). The signal of each electrode was referenced to a contact placed on the surface of the granulation tissue of either V1 or V4 chambers. Electrodes were connected to the recording system via a preamplifier (HS-36, Neuralynx) and the raw signal was collected with a 24-bit resolution at a sampling rate of 32 kHz. To obtain the LFP, the raw signal was band-pass filtered offline at 0.5-300 Hz and downsampled to 1017 Hz. To acquire the envelope of multiunit activity (MUA_E_), a Butterworth (3^rd^ order, 0.6-9 kHz) was applied on the raw signals and rectified. The envelope of the resulting signal was computed by a low-pass filtering (Butterworth 5^rd^ order, <300 Hz) (45). This was then downsampled to 1017 Hz.

### Bipolar re-referencing

Bipolar re-referencing was performed to remove the common signal on adjacent contacts and yield a local LFP. The LFP signal was re-referenced by subtracting the signal recorded at two neighboring channels. Specifically, the signal on channel *i* was acquired by obtaining the difference signal of channel *i*+1 and channel *i*-1.

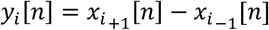

Where Y_i_ [n] denotes the signal of the re-referenced channel *i* which was located between the *i*+1 and *i*-1 electrodes.

### Ripple detection

The LFP re-referenced signal was band-pass filtered between (80-250 Hz, zero-lag linear phase FIR filter), squared and z-scored normalized. The mean and standard deviation (SD) was computed across the entire experimental duration to find the threshold for ripple event detection. Events from the normalized signal which exceeded 3.5 SD were selected as candidate ripples. Inter-ripple interval periods of less than 5 ms were merged. In addition, ripples of less than 30 ms duration and more than 100 ms were excluded. Finally, candidate ripples which exceeded 5 SD were considered to be a ripple (12, 46, 47). To avoid false ripple detection, we added an additional constraint. After detecting ripples by the algorithm, we calculated the spectrogram of ripples using a Hanning taper window. There were some false detected ripples with peak frequency power of less than 80 or above 200 Hz. Those falsely detected ripples were excluded from further analysis.

### Current source density (CSD) analysis

We used the inverse CSD (iCSD) toolbox to compute CSDs. CSD analysis was calculated by applying the spline method (48). LFPs were averaged over trial repetitions. Then, the iCSD, which is the second spatial derivative voltage (Ф) of LFPs, can be approximated using the following formula:

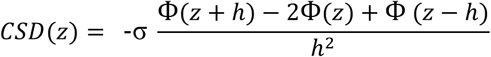

Where *z* is the depth at which the CSD is estimated, and *h* is the space between two electrodes (150µm). With this toolbox we used a Spline fitting method to interpolate Ф smoothly between electrode contacts. In our computation we assumed a tissue conductivity (σ) and a cortical column radius of 0.4 s/m and 500µm respectively (49, 50).

### Response latency analysis

details are given in supplementary materials.

### Laminar alignment

The CSD was used for alignment of probe contacts to the cortical layers with reference to layer IVc. Previous studies (51, 52) established that an early sink in the CSD profile corresponds to afferents terminating in layer IVc, with associated current inflow. First, we computed the CSD across all stimulus presentations of each session (Supplementary figure S5A) and visually determined the contact that featured the early sink (35-55 ms after stimulus onset; Supplementary figure S5B). Additionally, we determined the channel/contact with the shortest latency stimulus-evoked response MUA_E_ response (details in supplementary materials and supplementary figure S5C, D). Using these criteria, we assigned the reference contact in layer IVc and signals from other contacts were assigned to supra-granular, granular and infra-granular layers depending on their distance from the reference contact. For area V1, channels at 0.25 mm above and 0.25 mm below the reference channel were labelled as granular (IV), channels at 0.25 mm to 1 mm above the reference were labeled as supra-granular (I, II, III) and channels below the reference channel at 0.25 mm to 0.75 mm were labeled as infra-granular (V, VI) (40, 53). For V4, contacts less than 0.1 mm above and below the reference contact were identified as granular, 0.1 to 1 mm above the reference contact were identified as supra-granular, and those 0.1 mm to 0.75 mm below the reference channels were labelled as infra-granular (Supplementary figure S5, C, D). Channels outside these ranges were excluded from further analyses.

### Cross-correlation

The temporal relationship between ripples that occurred in V1 and V4 was explored using cross-correlation analysis. The cross-correlation (CC) was performed using the xcorr function in Matlab, according to:

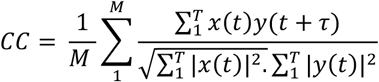

Where *x* and *y* are binary vectors representing ripple occurrence in area V1 and V4. For each trial, we made a vector of zeros for each area with ∼1 ms resolution (in line with the LFP sampling frequency).

If a ripple occurred, we replaced the content of that time point with a value of 1 and convolved it with a 50 ms Gaussian window. M and T denotes the number of trials and discrete time bins respectively and t represents the time lag. The area under the cross-correlation curve (AUC) was used to examine the synchrony of ripple occurrence between V1 and V4.

### Time-frequency spectral modulation

We estimated the power of the bipolar re-referenced LFP using the Chronux toolbox (54). Before Fourier transformation, the re-referenced LFP signal was mean-subtracted and multiplied with a Hanning taper. To compute spectrograms, short Fourier transformation was performed on a 200 ms length sliding window over the data with a 5 ms overlapping step. The spectrogram was corrected by the baseline spectrum to remove the 1/*f* effect.

### Number of ripples in behavioral epochs and peak frequency of ripples

Ripples were detected in three periods in the attention task, namely (i) pre-cue, (ii) post-cue and (iii) sustained activity after stimulus onset. To quantify the ripple rate for a particular task epoch, the sum of ripples detected for a given epoch was divided by the product of trial number and the epoch duration for a given recording session (sum of ripples / (number of trials × epoch duration)). Ripple rate was quantified separately for each session for all contacts combinations. Where relevant, statistics was performed over sessions, i.e. each session contributed a single ripple rate for a specific combination of conditions (e.g. pre-cue; or e.g. during the sustained period: attend RF AND small stimulus AND narrow focus of attention; etc.). The peak frequency within a SWR was calculated in a 200 ms window centered on SWR events with a Hanning window. A Kruskal Wallis test was used to determine whether peak frequencies differed between task epochs.

### Comodulogram

The Comodulogram was estimated to evaluate power-power coupling of ripples within and between V1 and V4. To estimate comodulograms, initially the wavelet-spectrogram was computed using Fast Fourier Transform (FFT) of bipolar re-referenced LFPs x(t) convolved with a FFT of complex Morlet-wavelets w(t,f). The output signal y(t,f) was acquired by the inverse FFT to estimate the time-frequency decomposition, where f denotes the desired center of the wavelet function. The wavelet’s central frequencies ranged from 3 to 250 Hz in 40 logarithmic spaced steps (55). Next, to compute the comodulogram, spectrograms of the two signals were computed as above. We refer to the first signal as trigger and to the second signal as target. For instance, if a ripple was detected in a trial in V1, the spectrogram of the ripple (trigger spectrogram) and the spectrogram of the exact time in V4’s trial (target spectrogram) was computed. Afterwards, in order to quantify the correlation of the two spectrograms, the Pearson correlation between the magnitude of power was calculated using the corr Matlab function (7, 56). The resulting correlation matrix was Fisher Z-transformed. This analysis was done across all contacts. The comudulogram was corrected by computing the comodulogram of random trials of the same conditions that did not contain ripples (matched for time during trial and exact condition combination). Finally, a t-test with false discovery rate correction (FDR) was conducted to find significant correlations (57). To estimate the comodulogram within an area, the same method was applied to calculate the power coupling between electrode contacts within and between layers.

### Spike rate during ripple time

Multi-unit spiking activity (MUA) was extracted from each recording channel by first bandpass filtering the raw data within the range of 600-9000Hz (sampled at 32756Hz). it was obtained by lowering a threshold progressively to obtain an average level of 100 Hz on each channel for the time relevant for our recording, namely the pre-cue, post-cue, and stimulus driven periods. Given that the 100Hz are averaged for periods prior to and after stimulus onset, stimulus driven activity exceeded the 100Hz average firing rate. Multi-unit spiking activity at the ripple time was computed within a 50 ms window centered at the ripple events across all attention conditions. To determine whether spiking activity varied with ripple events, we compared firing rate during ripples, with firing rates from the same number of trials where no ripples occurred, using matched time points within trials. For example, if a ripple occurred 412 ms after stimulus onset in one trial, we used the same time period in a condition/stimulus matched trial where no ripple occurred.

### Linear mixed-effects model

We used Matlab’s fitlme function to perform a linear mixed effect analysis of the relation between reaction times or firing rate with task parameters including ripple occurrence (yes/no), attention to receptive field (RF/away), focus of attention (narrow/wide) and size of stimulus (large/small) across all trials of the monkeys. Reaction times of both monkeys were concatenated (i.e. pooled across sessions) after z-score normalization for each session. Attention to the RF, size of stimulus, focus of attention and ripple occurrence were entered as fixed effects into the model as categorical variables, while different intercepts for each recording session were entered as random effects. Fixed effect variables were defined as ‘categorical’ in the model. Reaction times or firing rates were modelled as a linear combination of polynomial basis function of attention to RF (*Att*), ripple (*R*) occurrence, focus of attention (*F*) and size of stimulus (*S*), where β is the polynomial coefficient.

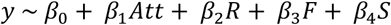

### Statistical testing

Kruskal-Wallis (KW) tests were used to compare ripple duration and ripple peak frequency between task epochs. Wilcoxon sign rank tests were used to determine whether there was a significant difference between ripple rate on different attention conditions and attention blocks. A one-way ANOVA was applied to compare ripple rates between layers. A repeated measure three-way ANOVA was performed on the interaction on task condition on ripple rate. Alpha values of p<0.05 were considered as statistically significant. For the comodulogram, we compared the power correlation using t-tests corrected for multiple comparison using false discovery rate (FDR) (57), using an α<0.05 as statistically significant.

## Supplementary information

### Supplementary Methods

Two adult male rhesus monkeys (10 and 12 kg, aged 8 and 9 years of age at the start of the experiments) were trained to comfortably sit in a plexiglass chair. Before training they had been implanted with a head holder device under sterile conditions. Two recording chambers were implanted over areas V1 and V4 under sterile conditions before electrophysiological recording. Details of surgical procedures, postsurgical management and analgesics have been described previously (58). All surgical and behavioral procedures conformed to the guidelines of UK Animals Scientific Procedures Act, European Communities Council Directive RL 2010/63/EC and the U.S. National Institutes of Health Guidelines for the Care and Use of Animals for Experimental Procedures. To promote behavioral motivation and performance, daily access to fluid was controlled during training and experimental period, using fluid control regimes that have minimal psychological impact and no measurable physiological impact on the animals (59). Animals were pair-housed in cages of >10m3 with play and foraging areas, swings, climbing areas, perches and ‘balconies’.

### Visual stimulation

Visual stimuli were generated on a personal computer and presented on a CRT monitor (Monkey 2: Dell Multiscan P1110, 21’’, mean luminance=90cd/m2; Monkey 1: Iiyama HM204DTA, 22’’, mean luminance=53cd/m2) with a refresh rate of 120 Hz, and a resolution of 1024×768 pixels. At the beginning of each session the monkeys’ eye position was calibrated. The eye position was monitored using an optical eye-tracker (ET-49, Thomas Recording, Giessen, Germany) with a sampling frequency of 200 Hz. Stimulus presentation and behavioral control was handled by Remote Cortex 5

### Basic Response Characterization (Receptive Field Mapping)

Prior to starting the attention paradigm, the classical receptive field was determined using a reverse-correlation paradigm, as described in detail previously (60). Briefly, dark squares with a size of 1°, 0.5° or 0.25° were used for mapping of receptive fields. Michelson contrast of the stimuli was 95-98 %. Stimuli were presented for 108 ms (monkey 1) or 125 ms (monkey 2) and covered a 9 × 12 grid of multiples the respective stimulus size.

RF maps were initially estimated online based on thresholded MUA spiking activity to determine the stimulus locations in the attention paradigm. Offline RF analysis was done based on local population activity (envelope multi-unit activity, MUA_E_), using a time window from 40-120 ms after RF mapping stimulus onset. MUA_E_ was computed as described in detail previously (45). Briefly, the higher frequency signal component (600-9000 Hz, (Butterworth, zero-phase digital filter of order 3)) was full-wave rectified, low-pass filtered at 200 Hz (Butterworth, zero-phase digital filter of order 3) and down sampled to a frequency of 1017 Hz.

We also determined orientation selectivity prior to the main recording sessions, and stimuli used were matched for the aggregate orientation preference obtained in a session where possible.

### Response Latency analysis

To estimate the latency of visual responses we used a method described previously (61). It makes two assumptions; (i) namely that the onset of the neuronal response has a Gaussian distribution across trials and (ii) that a fraction of response modulations dissipates exponentially after reaching the peak magnitude. These assumptions yield a function *f(t)* which consists of an ex-gaussian and a cumulative Gaussian function.

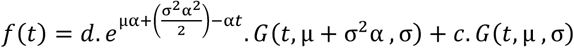

The parameters of µ and σ are the mean and standard deviation of the cumulative Gaussian, which are determined by onset response time. The parameter α is the dissipating rate and *c* and *d* act as weighting factors for the response magnitude and dissipation terms. For our data, we acquired 150 ms of MUA_E_ signal after stimulus onset, which was z-scored to baseline activity (before stimulus onset). The function *f(t)* was fit to the z-scored activity. The latency of visual responses (lat_33_) was determined as a point in time where the fitted function reached 33% of its maximum (45, 61).

### Ripple rate and RF overlap

To assess whether V1 and V4 RF overlap, and therefore stimulus placement, affected the ripple rate, we classified recording sessions into two categories. Namely, where V1 RFs were completely or almost completely covered by all of the V4 RFs (these are labelled as ‘fully overlapped’, examples in Supplementary figure S7) and sessions where the overlap was less complete (‘partial overlap’, example in Supplementary figure S7). Then, we quantified ripple rate for sessions with full and partial RF overlap. A one-way ANOVA revealed that there was no difference in ripple rate between sessions with full and partial overlap. This was the case for both V1 and V4 (F (1,732) = 1.57, p = 0.21, pooled data and Supplementary figure S8 panel A). Furthermore, we explored whether ripple rate modulation by stimulus size would change depending on RF overlap between V1 and V4. The reasoning here is that if small stimuli are centered on V1 RFs, which do not fully overlap with V4 RFs, V4 RFs would not, or only partially be driven by the small stimulus. This would not be the case if V1 and V4 have full overlap.

We explored the ripple rate among the fully overlapped sessions (n=18) in different task conditions. Corroborating the results reported in the main manuscript (where data were pooled across all session), small stimuli elicited a higher ripple than large stimuli in both areas. For these 18 recordings ripple rate for small and large stimuli were 0.15 Hz and 0.09 Hz in V1. In V4, rates were 0.11 Hz and 0.05 Hz for small and large stimuli (V1: Z = -4.2, p <0.001. V4: Z = -7.01, p<0.001, Wilcoxon’s signed rank test, Supplementary figure S8 panel B). In addition, attention to RF enhanced ripple rate when compared to attend away conditions among these sessions. Attend RF and away conditions in V1 triggered 0.13 Hz and 0.10 Hz respectively. In V4 attend RF conditions triggered ripple rates of 0.12 Hz while attend away conditions elicited 0.05 Hz (V1: Z = 2.08, p = 0.03. V4: Z = 8.11, p <0.001, Supplementary figure S8 panel C). We conducted a 3-way repeated measures ANOVA, to determine main effects of stimuli, attentional location and attentional focus, and possible interactions on ripple rate. ANOVA revealed that stimulus size (p=0.005) and attention location (p=0.02) had main effect on ripple rate frequency in V1 data. There was a significant attention location*size interaction (p=0.04) and attention location*attentional focus interaction (p=0.02) in V1. The 3-way ANOVA in V4 showed that stimulus size (p=0.0009) and attention location (p<0.001) significantly increased ripple rate. We found a stimulus size and attention location interaction on ripple rate (p=0.0003) in V4.

**Fig. S1.**
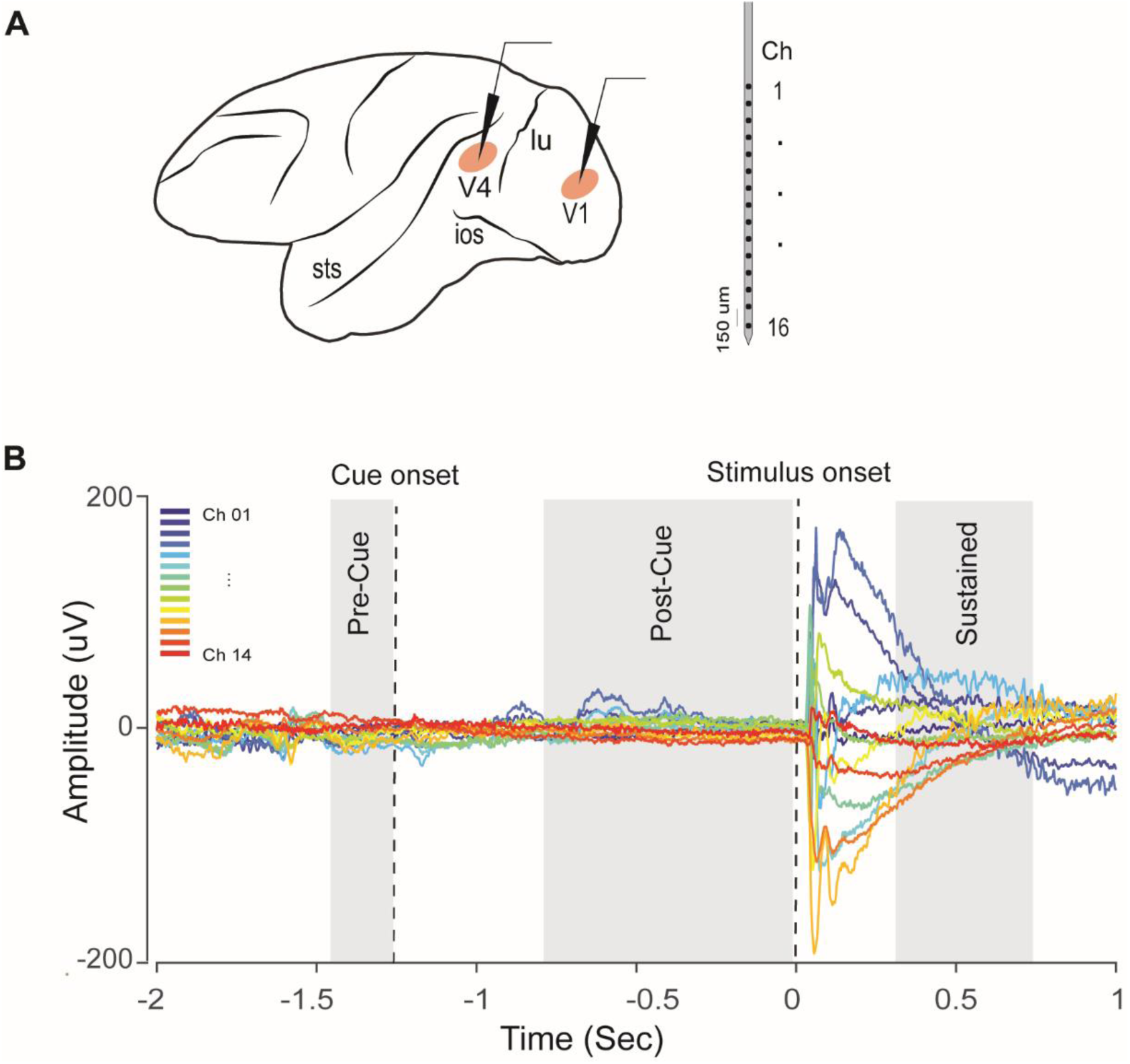
Laminar probe and event related potential (ERP). A) Schematic representation of 16-channel laminar probe inserted into V1 and V4 for collecting of spikes and LFPs. B) Traces of ERPs acquired by the probe and task epochs used for detection of ripples. Each color depicts average of a bipolar re-referenced LFP acquired by a contact. Time of epochs aligned to stimulus onset (time 0). Black dashed lines denote the onset and offset of cue and stimulus.

**Fig. S2.**
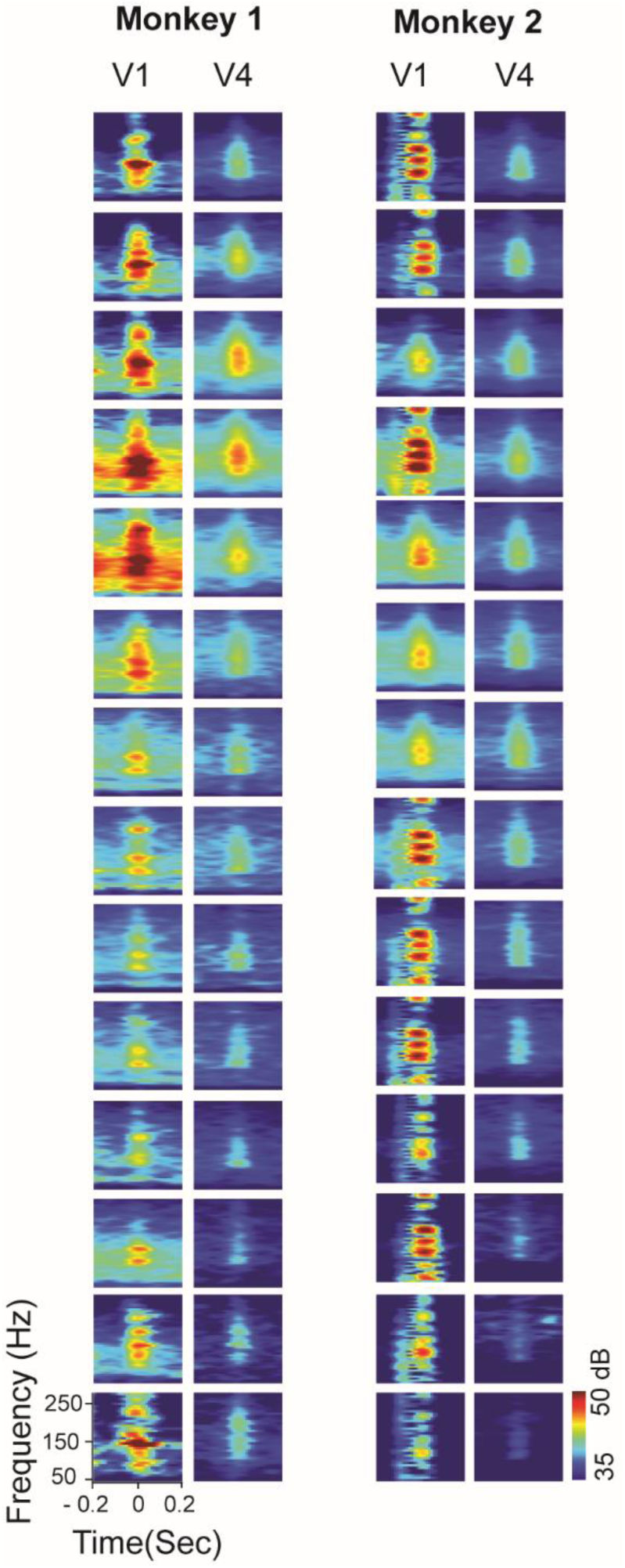
Depth profile of spectrogram of ripples across electrode contacts. Left panel indicates grand average of spectrogram of ripples identified on bipolar-referenced LFPs acquired by laminar probes in Monkey 1 in V1 and V4. Right panel shows data for monkey 2.

**Fig. S3.**
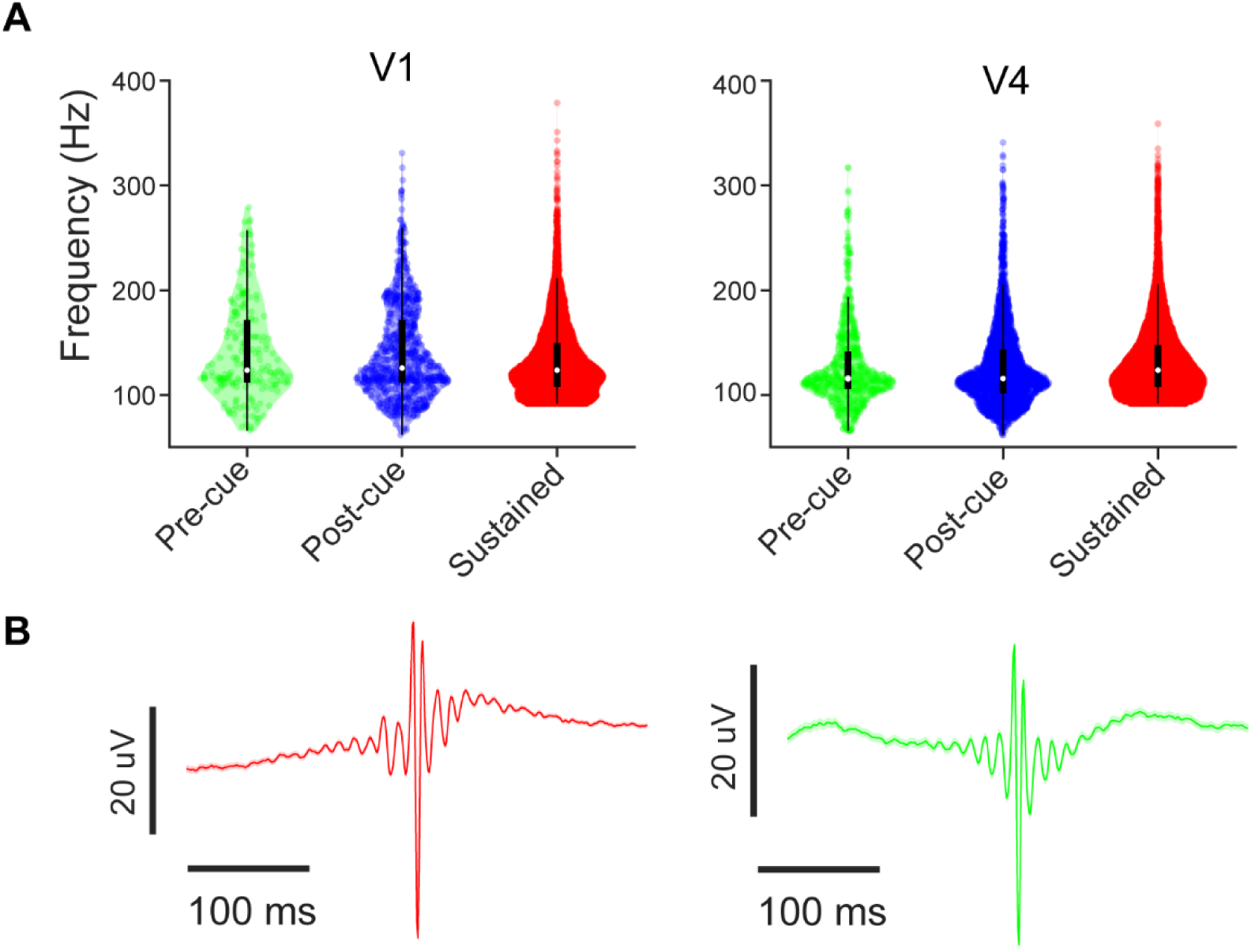
Ripple’s peak frequency and mean peri-ripple field potentials in V1 and V4. A) Peak frequency of ripples among pre-cue, post-cue and sustained intervals. Ripples peak frequency in V1 and V4 was calculated from a 200 ms window centered at ripple time. Ripples in the sustained period showed lower peak frequency compared to pre-cue and post-cue conditions in V1 as well as V4. B) Group-averaged ripple-triggered wide-band traces (<300 Hz) for V1 and V4.

**Fig. S4.**
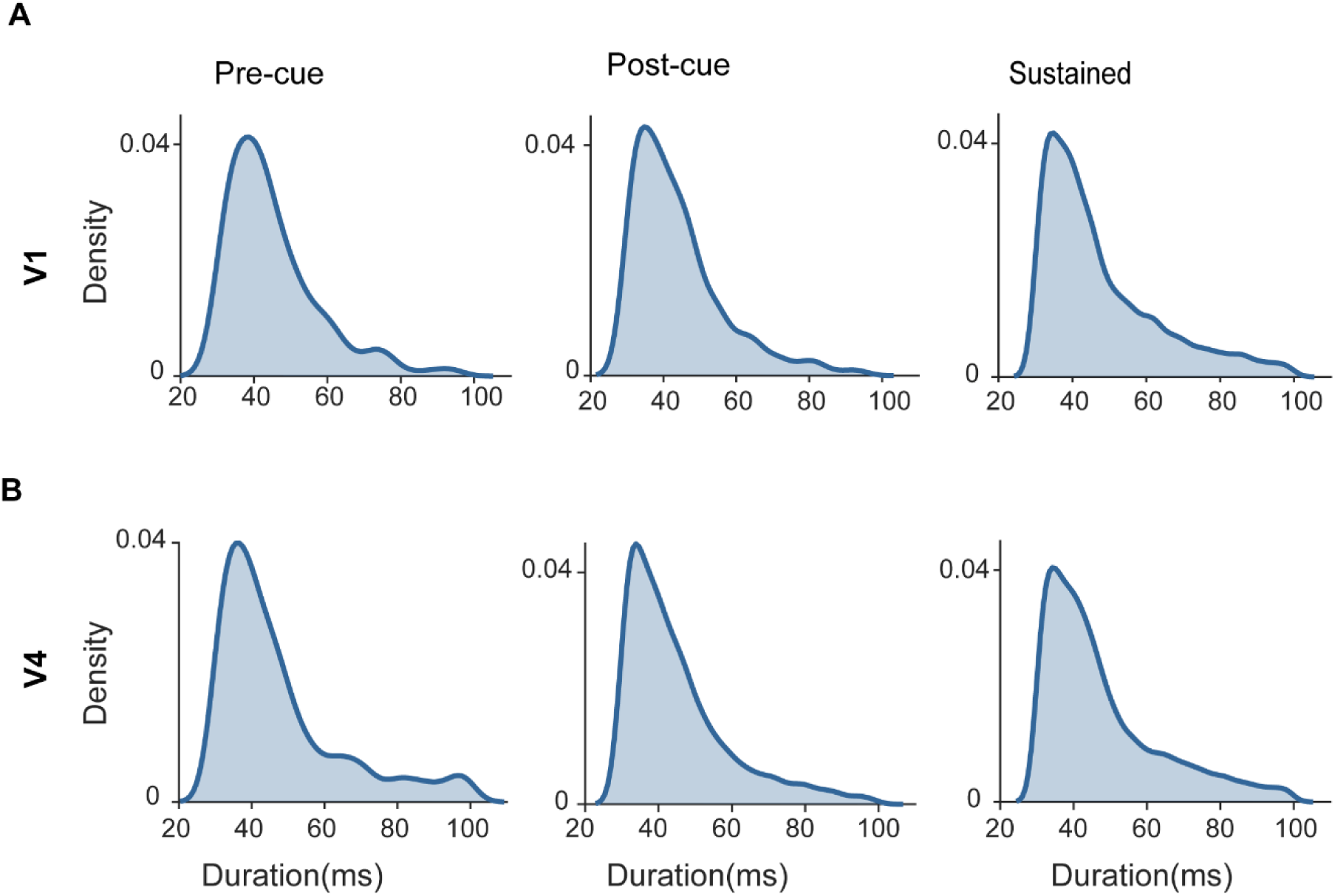
Kernel density of ripple duration during different task epochs for V1 and V4. Ripple duration was longer during the sustained period than post and peri cue periods in V1 and V4.

**Fig. S5.**
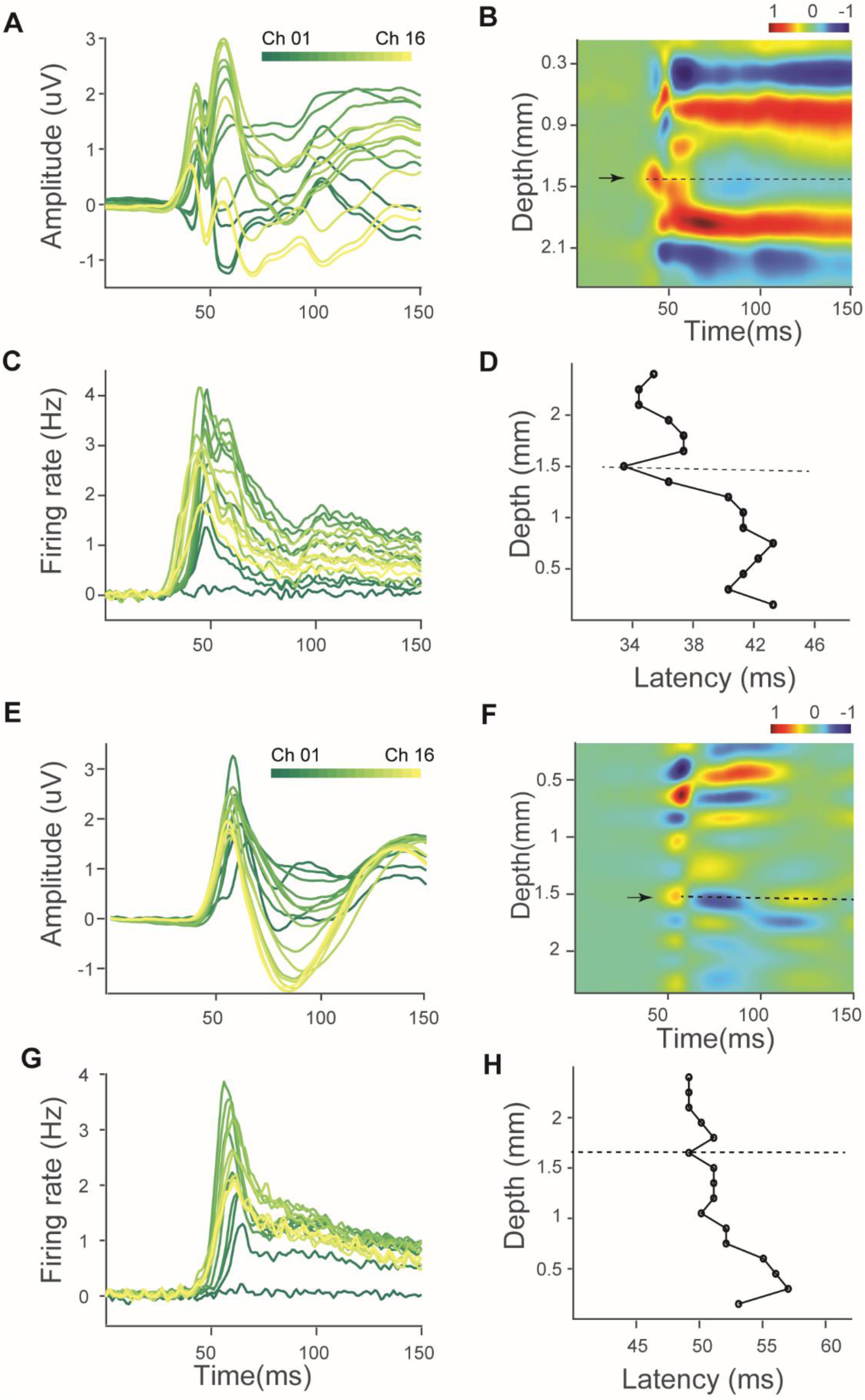
Samples of laminar alignment in V1 (A to D) and V4 (E to H). A, E) LFPs z-scored to the baseline activity. Different colors denote activity acquired by different contacts. C, G) MUAE z-scored to baseline activity used to compute latency of each contact. B, F) CSD z-scored to baseline. Black arrow represents the first input to the cortex (sink region). The dotted line is the depth defined for alignment of the layers. D, H) are response latency indices across depth.

**Fig. S6.**
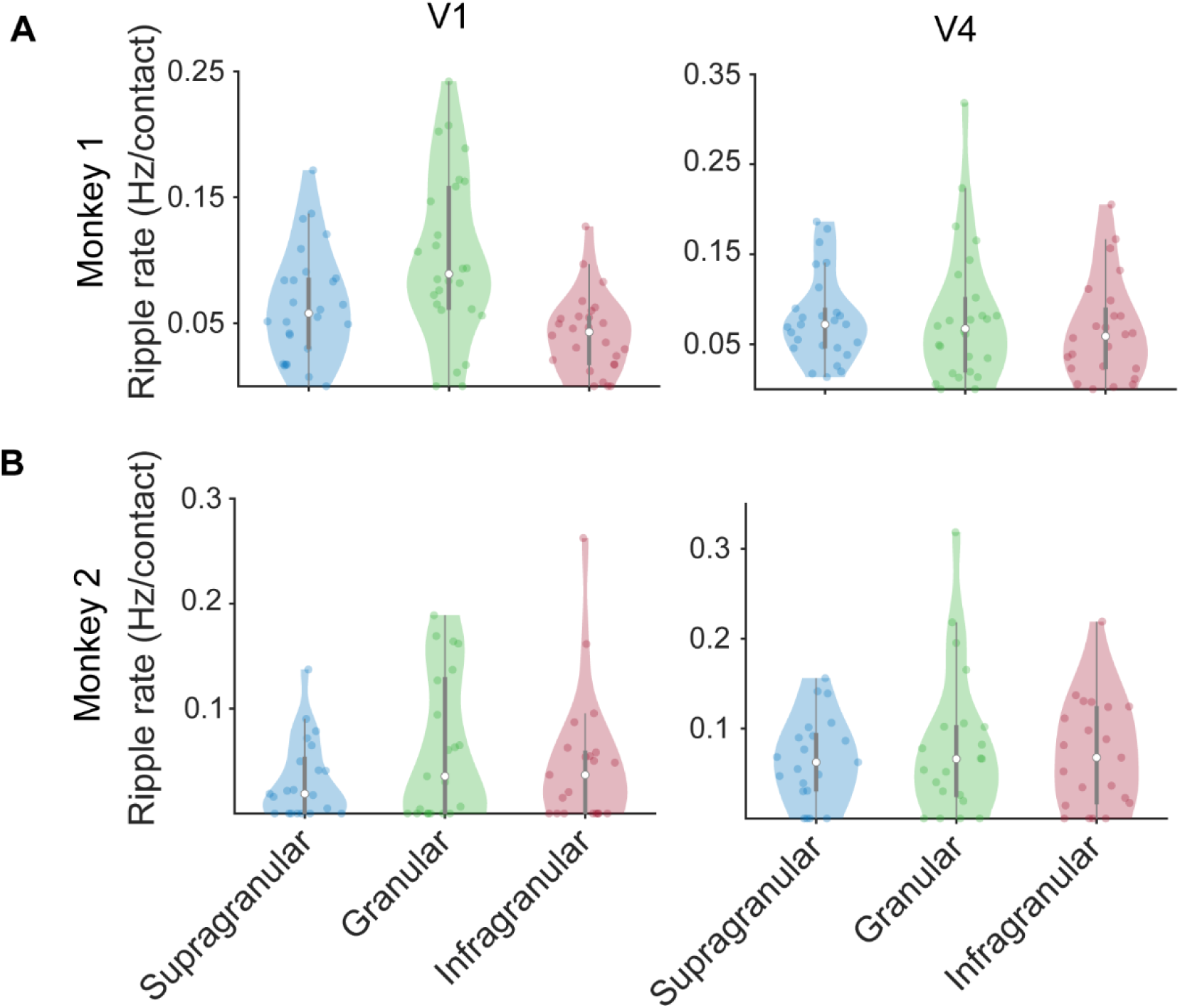
Ripple rate across cortical depth. A) Left panel illustrates ripple rate for V1, and right panel represents ripple rate for V4 layers (per contact) for supragranular, granular and Infragranular layers in monkey 1. Ripple rate was larger for granular layer in V1 compared with infragranular layer but did not differ otherwise between compartment comparisons. B) Same as panel A for monkey 2.

**Fig. S7.**
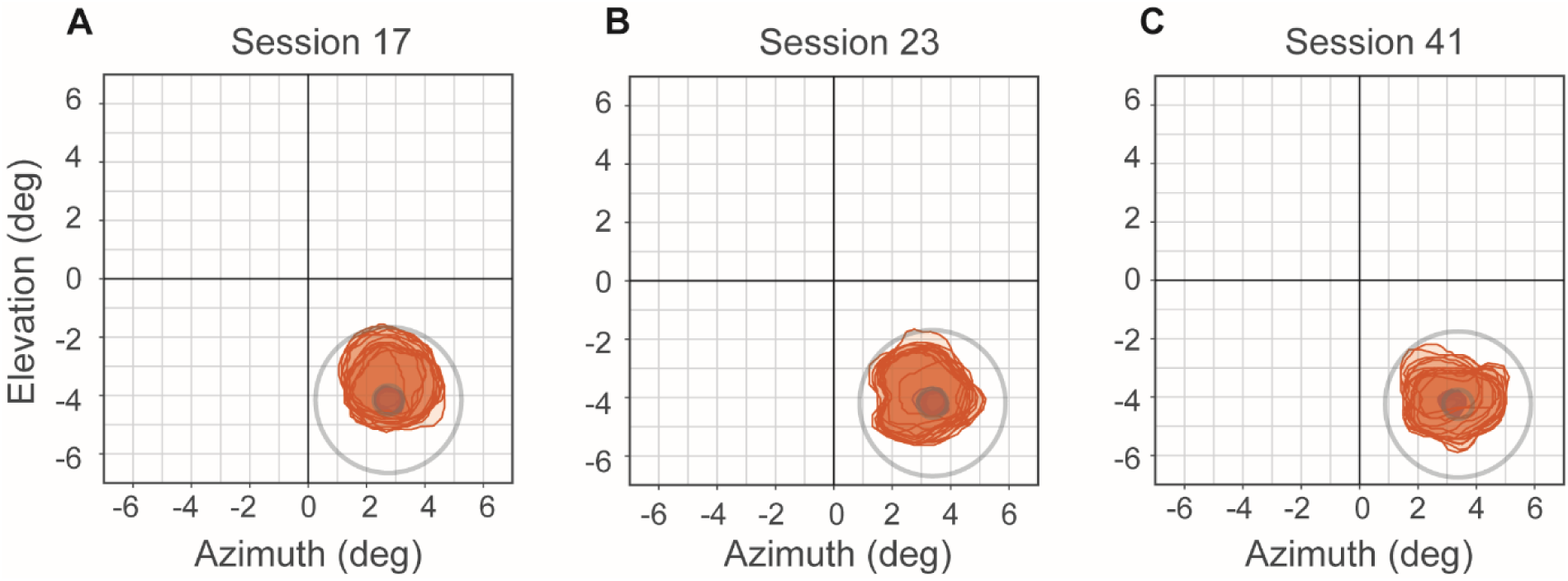
Illustration of three sample recordings with fully overlapping V1 and V4 receptive fields. Panels depict V1 and V4 RFs obtained from MUAE activity for each electrode contact using 0.25**°**, 0.5**°** and 1**°** sized stimuli. RF contours are relative to fixation point (point 0,0, units are in degrees of visual angle). Blue colors are V1 RFs, covered by V4 RFs in red.

**Fig. S8.**
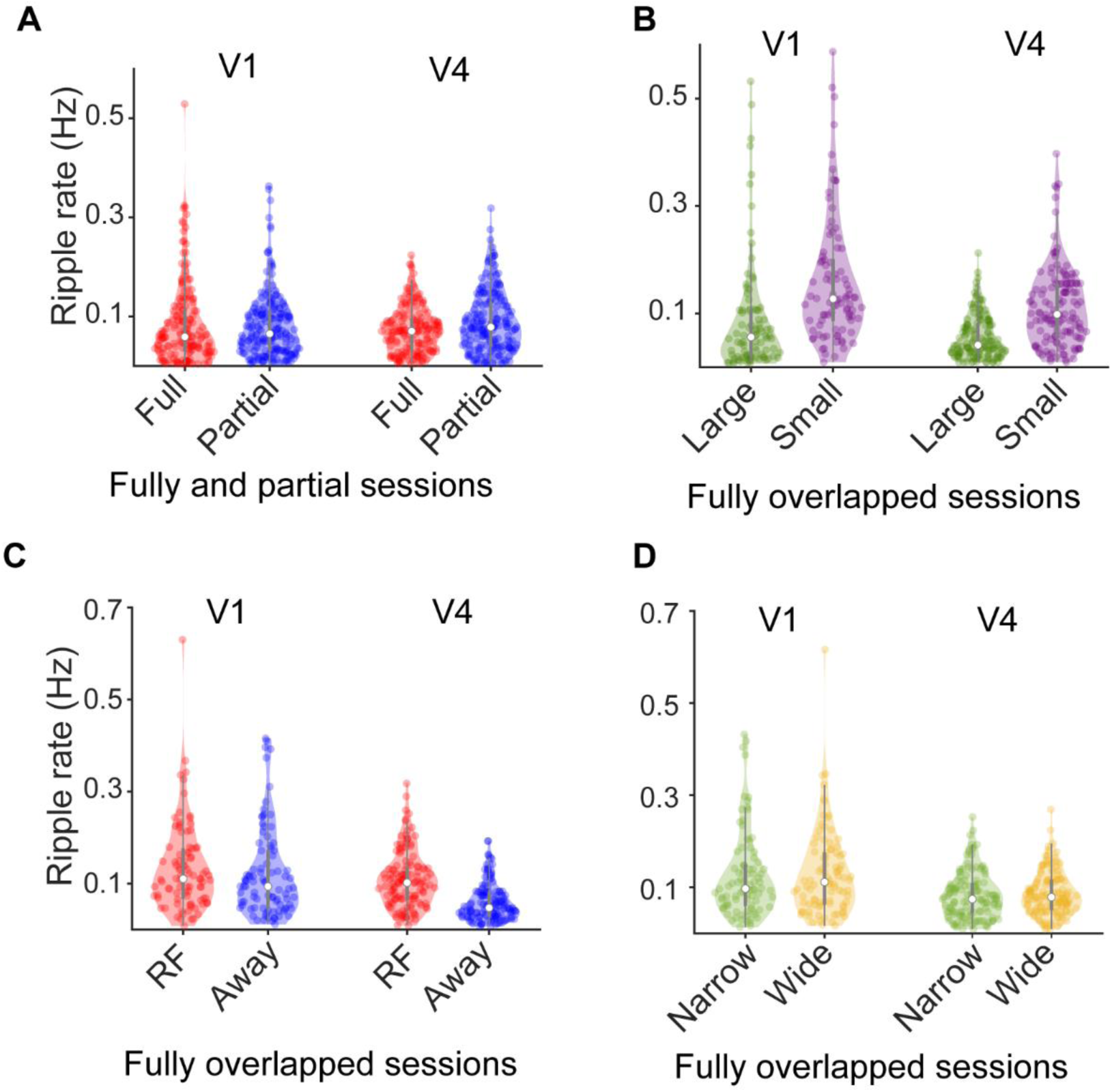
Ripple rate with respect to receptive field (RF) overlap between V1 and V4. A) Comparison of ripple rate as function of overlapping of V1 and V4 RFs. Red and blue colors represent ripple rate quantified between recordings where RFs of V1 were completely (full) or partially (partial) overlapping with V4 RFs. B) Effect of stimulus size on ripple rates for recordings with full overlapping V1 and V4 RFs. C) Ripple rate in attend to RF/away conditions for session with full RF overlap. Similar to the results reported in the main manuscript, sessions with full V1-V4 RF overlap, show that small stimuli and attend RF elicited higher ripple than large and attend away condition. This effect was consistent in V1 and V4. D) Ripple rate computed on the narrow and wide blocks among the sessions with fully overlapping of the RFs in V1 and V4.

**Fig. S9.**
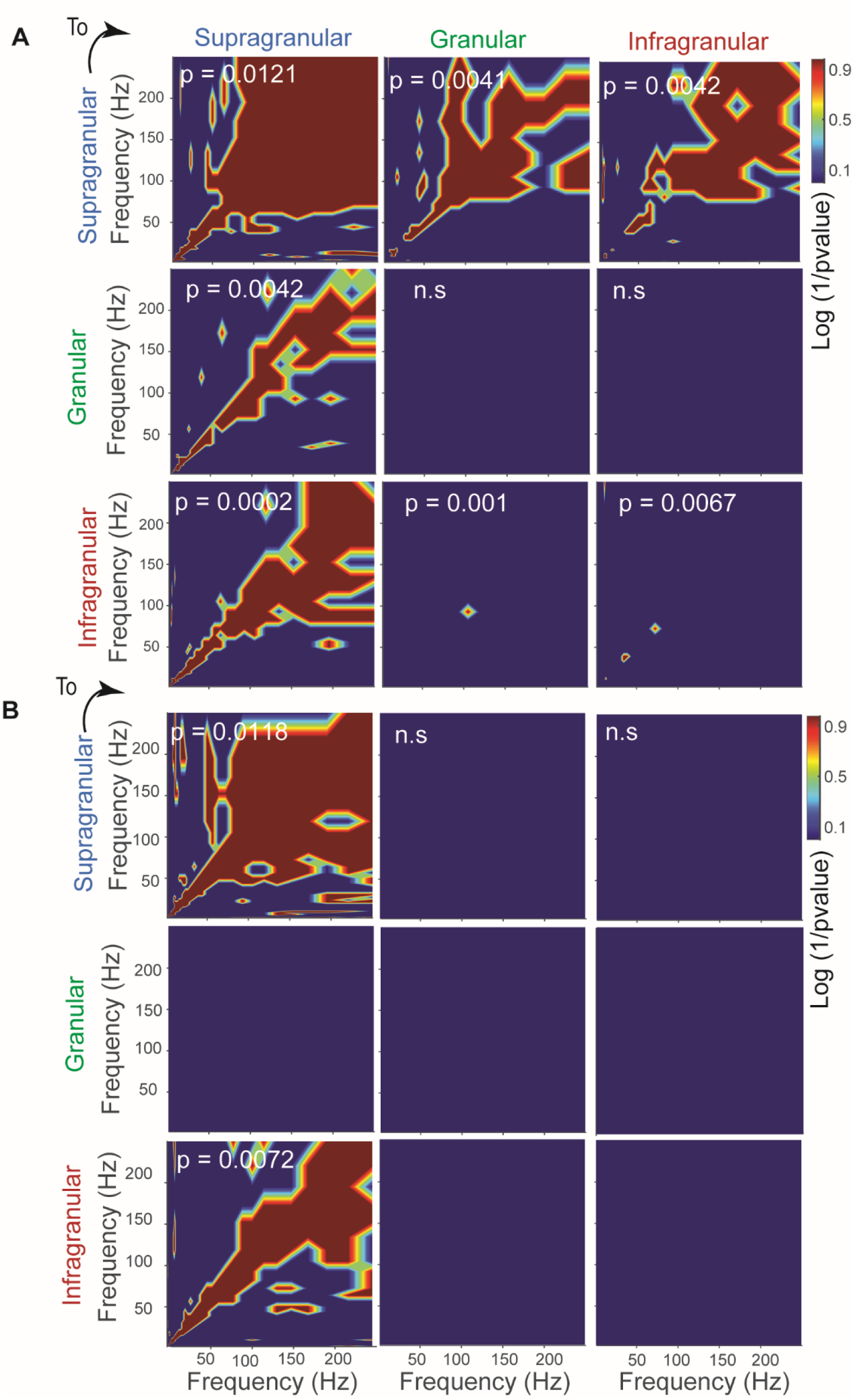
Within layer power correlation. A) Corrected comodulogram between supragranular, granular and infragranular in V1. Each panel shows distribution of p-values, log10(1/p-values), between comodulogram of layers where a specific layer served as a trigger. The arrow at top left denotes the direction of the trigger region and the respective response. Red areas show frequency bands that showed significant power coupling (t-test with FDR correction). B) Comodulogram of ripple in V4 between different layers. Same as (A) but for area V4.

**Fig. S10.**
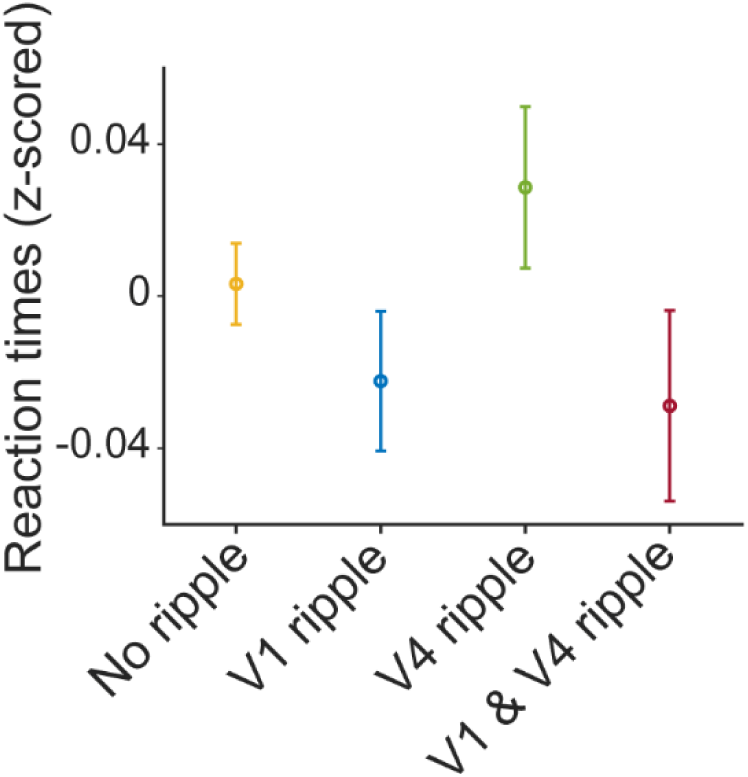
Reaction time across trials with and without ripples in V1 and V4. Mean and SEM of z-scored reaction time during trials when no ripples occurred during the sustained period, when ripples occurred only in V1 during the sustained period, when they occurred only in V4 and when the occurred in both areas, V1 and V4.

**Table S1.**
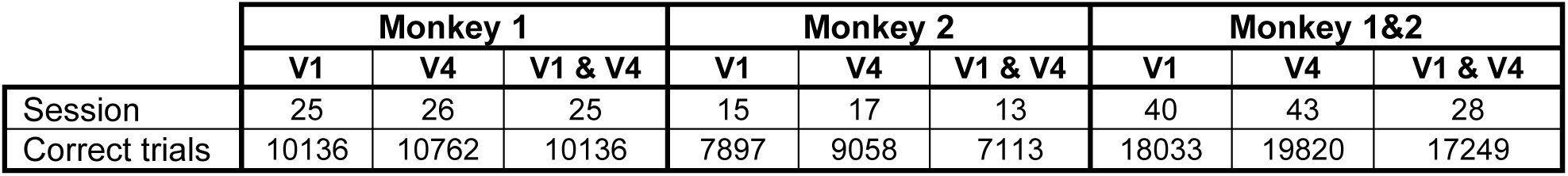
Recording sessions and trials. Number of sessions for data recorded in V1 and V4. The columns V1& V4 report the number of sessions and trials where data were recorded simultaneously in both V1 and V4.

**Table S2.** Three-Factor (Stimulus, Location of Attention, Attentional Focus) Analysis of Variance with Repeated Measures to assess modulation of V1 ripple rate (during the sustained period)

|  | **SS** | **df** | **MS** | **F** | **P** |
| --- | --- | --- | --- | --- | --- |
| Between-Sessions | 0.419 | 39 |  |  |  |
| Within-Sessions | 1.594 | 280 |  |  |  |
| **Stimulus** | 0.560 | 1 | 0.560 | 31.233 | <0.001 |
| error (stimulus) | 0.530 | 39 | 0.014 |  |  |
| **Attention** | 0.016 | 1 | 0.016 | 6.564 | 0.014 |
| error (attention) | 0.096 | 39 | 0.002 |  |  |
| **Focus** | 0.000 | 1 | 0.000 | 0.016 | 0.902 |
| error(focus) | 0.075 | 39 | 0.002 |  |  |
| **Stimulus\* Attention** | 0.006 | 1 | 0.006 | 4.796 | 0.035 |
| Error(stimulus*attention) | 0.176 | 39 | 0.004 |  |  |
| **Stimulus\*Focus** | 0.036 | 1 | 0.036 | 25.765 | <0.001 |
| Error(stimulus*focus) | 0.040 | 39 | 0.001 |  |  |
| **Attention\*Focus** | 0.012 | 1 | 0.012 | 7.562 | 0.009 |
| Error(attention*focus) | 0.043 | 39 | 0.001 |  |  |
| **Stimulus\*Attention\*Focus** | 0.006 | 1 | 0.006 | 0.150 | 0.7476 |
| Error | 0.055 | 39 | 0.001 |  |  |
| **Total** | 1.978 | 639 |  |  |  |

3-way repeated measures ANOVA to determine whether ripple rate in area V1 during the sustained period depended on stimulus size (small/large), locus of attention (attend RF/away) and attentional focus size (narrow/wide). Table denotes sum of squares (SS), degrees of freedom (df), mean squares (MS), F-value (F), and p-value (P). Interaction terms are indicated by \* sign.

**Table S3.** Three-Factor (Stimulus, Location of Attention, Attentional Focus) Analysis of Variance with Repeated Measures to assess modulation of V4 ripple rate (during the sustained period)

|  | **SS** | **df** | **MS** | **F** | **P** |
| --- | --- | --- | --- | --- | --- |
| Between-Sessions | 0.711 | 42 |  |  |  |
| Within-Sessions | 2.288 | 301 |  |  |  |
| **Stimulus** | 0.530 | 1 | 0.530 | 31.233 | <0.001 |
| error (stimulus) | 0.712 | 42 | 0.017 |  |  |
| **Attention** | 0.378 | 1 | 0.378 | 110.428 | <0.001 |
| error (attention) | 0.144 | 42 | 0.003 |  |  |
| **Focus** | 0.011 | 1 | 0.011 | 8.681 | 0.005 |
| error(focus) | 0.052 | 42 | 0.001 |  |  |
| **Stimulus\* Attention** | 0.142 | 1 | 0.142 | 33.830 | <0.001 |
| Error(stimulus*attention) | 0.176 | 42 | 0.004 |  |  |
| **Stimulus\*Focus** | 0.008 | 1 | 0.008 | 7.871 | 0.008 |
| Error(stimulus*focus) | 0.040 | 42 | 0.001 |  |  |
| **Attention\*Focus** | 0.007 | 1 | 0.007 | 6.810 | 0.013 |
| Error(attention*focus) | 0.043 | 42 | 0.001 |  |  |
| **Stimulus\*Attention\*Focus** | 0.002 | 1 | 0.000 | 0.048 | 0.8282 |
| Error | 0.036 | 42 | 0.001 |  |  |
| **Total** | 2.991 | 687 |  |  |  |

3-way repeated measures ANOVA to determine whether ripple rate in area V4 during the sustained period depended on stimulus size (small/large), locus of attention (attend RF/away) and attentional focus size (narrow/wide). Table denotes sum of squares (SS), degrees of freedom (df), mean squares (MS), F-value (F), and p-value (P). Interaction terms are indicated by \* sign.

**Table S4.** Multiple linear mixed effect model results to predict firing rates in V1 from attention, ripples, attentional focus and stimulus size.

| Predictors (V1) | $\beta$ | T | p-values |
| --- | --- | --- | --- |
| Ripple (0) | -11.4 | -15.5 | < 0.0001*** |
| Attention (1) | 0.36 | 0.49 | 0.6 |
| Focus (1) | -2.3 | -3.1 | 0.001*** |
| Size (1) | -19.3 | -26.3 | < 0.001 *** |
| Ripple (0) * attention (1) | 1.2 | 1.7 | 0.08 |
| Ripple (0) * Focus (1) | -0.9 | -1.2 | 0.2 |
| Attention (1) * Focus (1) | 0.15 | 0.2 | 0.8 |
| Ripple (0) * size (1) | -0.9 | -1.2 | 0.2 |
| Attention (1) * size (1) | 0.01 | 0.01 | 0.9 |
| Focus (1) * size (1) | -0.37 | -0.5 | 0.6 |
| Ripple (0) * attention (1) * Focus (1) | -0.29 | -0.4 | 0.6 |
| Ripple (0) * attention (1) * size (1) | 0.4 | 0.5 | 0.5 |
| Ripple (0) * Focus (1) * size (1) | -0.7 | -0.9 | 0.3 |
| Attention (1) * Focus (1) * size (1) | 1.1 | 1.5 | 0.1 |
| Ripple (0) * Attention (1) * Focus (1) * size (1) | -0.49 | -0.6 | 0.4 |

Linear regression model to predict V1 firing rate from occurrence of ripple, attention to RF vs away, size of focus of attention (narrow/wide), and size of stimulus (small/large). The digits in front the predictors represent the categories assigned to fixed effects into the model. Ripple (0 is without and 1 with ripple), attention (1 attention to RF 2 and away), focus (1 narrow and 2 wide focus of attention) and size (1 large and 2 small stimuli). The column of $\beta$ reports the coefficients of each predictor estimated by model and column T is the corresponding t-statistic. Asterisks indicate significance.

**Table S5.** Mean and SEM of firing rate across trials with and without ripple and during different attention conditions in V1.

| Ripple | Attention | Mean ± SEM of firing rate |
| --- | --- | --- |
| No | RF | 144.67 ± 1.2 |
| No | away | 140.68 ± 1.4 |
| Yes | RF | 164.96 ± 1.3 |
| Yes | Away | 165.28 ± 1.5 |

Column 1 indicates ripple occurrence in trials (with: yes; without: no ripple). Column 2 attention indicates attend to RF/away from RF condition.

**Table S6.** Multiple linear mixed effect model in V4.

| Predictors (V4) | $\beta$ | T | p-values |
| --- | --- | --- | --- |
| Ripple (0) | -11.8 | -21 | < 0.001 *** |
| Attention (1) | 6.3 | 11.3 | < 0.001 *** |
| Focus (1) | 0.4 | 0.8 | 0.3 |
| Size (1) | -7.1 | -12.6 | < 0.001 *** |
| Ripple (0) * attention (1) | 0.7 | 1.3 | 0.1 |
| Ripple (0) * Focus (1) | 0.9 | 1.7 | 0.08 |
| Attention (1) * Focus (1) | 0.1 | 0.3 | 0.7 |
| Ripple (0) * size (1) | -1.2 | -2.1 | 0.03* |
| Attention (1) * size (1) | -2.3 | -4.1 | < 0.001 *** |
| Focus (1) * size (1) | 1.6 | 2.9 | 0.003 |
| Ripple (0) * attention (1) * focus (1) | -0.4 | -0.7 | 0.4 |
| Ripple (0) * attention (1) * size (1) | 0.6 | 1.1 | 0.2 |
| Ripple (0) * focus (1) * size (1) | -0.1 | -0.2 | 0.8 |
| Attention (1) * Focus (1) * size (1) | -0.8 | -1.5 | 0.1 |
| Ripple (0) * Attention (1) * Focus (1) * size (1) | -0.3 | -0.5 | 0.5 |

Using a linear regression model to predict V4 firing rate from occurrence of ripple, attention to RF vs away, size of focus of attention (narrow/wide), and size of stimulus (small/large). The digits in front the predictors represent the categories assigned to fixed effects into the model. Ripple (0 is without and 1 with ripple), attention (1 attention to RF 2 and away), focus (1 narrow and 2 wide focus of attention) and size (1 large and 2 small stimuli). The column of $\beta$ reports the coefficients of each predictor estimated by model and column T is corresponding t-statistic. Asterisks indicate significance.

**Table S7.**
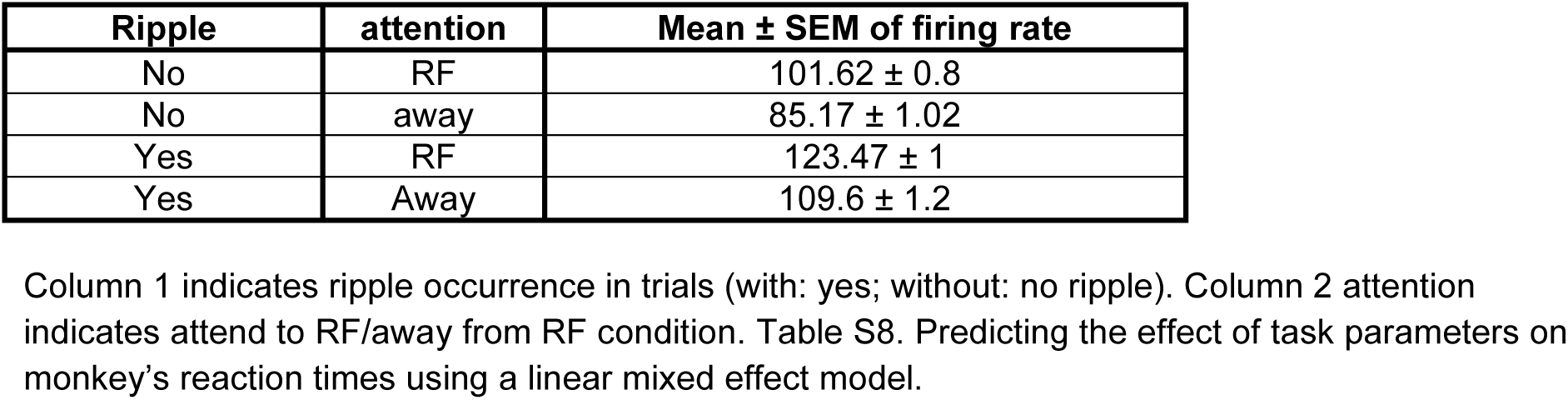
Mean and SEM of firing rate across trials with and without ripple during different attention conditions in V4.

| Ripple | attention | Mean ± SEM of firing rate |
| --- | --- | --- |
| No | RF | 101.62 ± 0.8 |
| No | away | 85.17 ± 1.02 |
| Yes | RF | 123.47 ± 1 |
| Yes | Away | 109.6 ± 1.2 |

**Table S8.** Predicting the effect of task parameters on monkey’s reaction times using a linear mixed effect model.

| Predictors | $\beta$ | T | p-values |
| --- | --- | --- | --- |
| Ripple (0) | -0.004 | -0.3 | 0.7 |
| Ripple (1) | -0.02 | -1.3 | 0.1 |
| Ripple (2) | 0.02 | 1.1 | 0.2 |
| Attention (1) | -0.01 | -1.4 | 0.1 |
| Focus (1) | 0.01 | 1.6 | 0.1 |
| Size (1) | -0.005 | -0.52 | 0.6 |
| Ripple (0) * attention (1) | -0.01 | -0.86 | 0.3 |
| Ripple (1) * attention (1) | -0.002 | -0.13 | 0.8 |
| Ripple (2) * attention (1) | -0.005 | -0.31 | 0.7 |
| Ripple (0) * Focus (1) | 0.006 | 0.47 | 0.6 |
| Ripple (1) * focus (1) | 0.01 | 0.70 | 0.4 |
| Ripple (2) * focus (1) | 0.01 | 0.65 | 0.5 |
| Attention (1) * focus (1) | 0.02 | 2.0 | 0.03 * |
| Ripple (0) * size (1) | -0.009 | -0.74 | 0.4 |
| Ripple (1) * size (1) | -0.02 | -1.5 | 0.1 |
| Ripple (2) * size (1) | -0.02 | -1.4 | 0.1 |
| Attention (1) * size (1) | -0.02 | -2.6 | 0.007 *** |
| Focus (1) * size (1) | -0.13 | -12 | < 0.001 *** |
| Ripple (0) * attention (1) * focus (1) | -0.01 | -0.9 | 0.3 |
| Ripple (1) * attention (1) * Focus (1) | 0.02 | 1.4 | 0.1 |
| Ripple (2) * attention (1) * focus (1) | -0.01 | -0.58 | 0.5 |
| Ripple (0) * attention (1) * size (1) | 0.004 | 0.35 | 0.7 |
| Ripple (1) * attention (1) * size (1) | -0.03 | -1.8 | 0.05 * |
| Ripple (2) * attention (1) * size (1) | 0.01 | 0.66 | 0.5 |
| Ripple (0) * focus (1) * size (1) | 0.02 | 2.2 | 0.02 ** |
| Ripple (1) * focus (1) * size (1) | 0.01 | 0.71 | 0.4 |
| Ripple (2) * focus (1) * size (1) | -0.008 | -0.46 | 0.6 |
| Attention (1) * focus (1) * size (1) | 0.01 | 0.96 | 0.3 |
| Ripple (0) * attention (1) * focus (1) * size (1) | 0.01 | 1.0 | 0.2 |
| Ripple (1) * attention (1) * focus (1) * size (1) | 0.003 | 0.1 | 0.8 |
| Ripple (2) * attention (1) * focus (1) * size (1) | 0.001 | 0.08 | 0.9 |

Results of the linear mixed effect model predicting reaction times from occurrence of ripple, attention to RF, stimulus size and narrow and wide attention. Numbers in the predictor's column indicate a category assigned to each variable. Ripple (0: no ripple, 1: ripple occurrence in V1, 2: ripple occurrence in V4, 3: ripple occurrence in V1&V4), attention (1: attention to RF, 2: attention away), focus (1: narrow and 2: wide focus of attention) and size (1: large and 2: small stimuli). Asterisks denote significance. B and T are the coefficient estimate and t-statistics.

**Table S9.** Mean and SEM of reaction time with respect to ripple occurrence.

| Ripple | Mean ± SEM of firing rate |
| --- | --- |
| No | 0.003 ± 0.01 |
| V1 | -0.022 ± 0.01 |
| V4 | 0.028 ± 0.02 |
| V1 & V4 | -0.028 ± 0.02 |

Z-scored reaction times are grouped by trials without ripple (no), ripple occurred in V1, V4 or both areas.

## Author Contributions

Jafar Doostmohammadi: Data acquisition, data analysis and analysis methods, data curation, manuscript writing, visualization.

Marc Alwin Gieselmann: Data acquisition, data analysis and analysis methods, data curation, manuscript review, visualization.

Jochem van Kempen: Data analysis, manuscript review.

Ali Yoonessi: Funding acquisition, manuscript review.

Reza Lashgari: Manuscript review.

Alexander Thiele: Conceptualization, data acquisition and analysis, manuscript writing and review, supervision, funding acquisition.

## Acknowledgments

Funded by the Wellcome Trust [093104] (JvK, AT), MRC [MR/P013031/1] (JvK, MAG, AT), MR/K013785/1 (MAG, AT), Tehran University of Medical Sciences [grant No. 97-02-87-38826] (JD Ph. D dissertation). We would like to thank Ali Ghazizadeh, Institute for Research in Fundamental Sciences, School of cognitive sciences, for intellectual support. We also thank the Newcastle CBC staff for excellent support.

## References

1. A. Bragin, J. Engel, C. L. Wilson, I. Fried, G. Buzsáki, High-frequency oscillations in human brain. Hippocampus 9, 137–142 (1999).

2. G. Buzsáki, Z. Horváth, R. Urioste, J. Hetke, K. Wise, High-frequency network oscillation in the hippocampus. Science 256, 1025–1027 (1992).

3. N. K. Logothetis, et al., Hippocampal-cortical interaction during periods of subcortical silence. Nature 491, 547–553 (2012).

4. Y. Nir, et al., Regional Slow Waves and Spindles in Human Sleep. Neuron 70, 153–169 (2011).

5. Y. Norman, et al., Hippocampal sharp-wave ripples linked to visual episodic recollection in humans. Science 365 (2019).

6. A. Ylinen, et al., Sharp wave-associated high-frequency oscillation (200 hz) in the intact hippocampus: Network and intracellular mechanisms. J. Neurosci. 15, 30–46 (1995).

7. D. Khodagholy, J. N. Gelinas, G. Buzsáki, Learning-enhanced coupling between ripple oscillations in association cortices and hippocampus. Science 358, 369–372 (2017).

8. G. Buzsáki, Hippocampal sharp wave-ripple: A cognitive biomarker for episodic memory and planning. Hippocampus 25, 1073–1188 (2015).

9. H. R. Joo, L. M. Frank, The hippocampal sharp wave–ripple in memory retrieval for immediate use and consolidation. Nat. Rev. Neurosci. 19, 744–757 (2018).

10. V. Ego-Stengel, M. A. Wilson, Disruption of ripple-associated hippocampal activity during rest impairs spatial learning in the rat. Hippocampus 20, 1–10 (2010).

11. D. J. Foster, M. A. Wilson, Reverse replay of behavioural sequences in hippocampal place cells during the awake state. Nature 440, 680–683 (2006).

12. G. Girardeau, K. Benchenane, S. I. Wiener, G. Buzsáki, M. B. Zugaro, Selective suppression of hippocampal ripples impairs spatial memory. Nat. Neurosci. 12, 1222–1223 (2009).

13. A. K. Lee, M. A. Wilson, Memory of sequential experience in the hippocampus during slow wave sleep. Neuron 36, 1183–1194 (2002).

14. B. P. Staresina, et al., Hierarchical nesting of slow oscillations, spindles and ripples in the human hippocampus during sleep. Nat. Neurosci. 18, 1679–1686 (2015).

15. M. A. Wilson, B. L. McNaughton, Reactivation of hippocampal ensemble memories during sleep. Science 265, 676–679 (1994).

16. D. Dupret, J. O’Neill, B. Pleydell-Bouverie, J. Csicsvari, The reorganization and reactivation of hippocampal maps predict spatial memory performance. Nat. Neurosci. 13, 995–1002 (2010).

17. B. E. Pfeiffer, D. J. Foster, Hippocampal place-cell sequences depict future paths to remembered goals. Nature 497, 74–79 (2013).

18. A. C. Singer, M. F. Carr, M. P. Karlsson, L. M. Frank, Hippocampal SWR Activity Predicts Correct Decisions during the Initial Learning of an Alternation Task. Neuron 77, 1163–1173 (2013).

19. G. Dragoi, S. Tonegawa, Preplay of future place cell sequences by hippocampal cellular assemblies. Nature 469, 397–401 (2011).

20. G. Buzsáki, L. Lai-Wo S. C. H. Vanderwolf, Cellular bases of hippocampal EEG in the behaving rat. Brain Res. Rev. 6, 139–171 (1983).

21. W. E. Skaggs, et al., EEG Sharp Waves and Sparse Ensemble Unit Activity in the Macaque Hippocampus. J. Neurophysiol. 98, 898–910 (2007).

22. N. Axmacher, C. E. Elger, J. Fell, Ripples in the medial temporal lobe are relevant for human memory consolidation. Brain 131, 1806–1817 (2008).

23. T. K. Leonard, et al., Sharp wave ripples during visual exploration in the primate hippocampus. J. Neurosci. 35, 14771–14782 (2015).

24. J. O’Neill, T. Senior, J. Csicsvari, Place-Selective Firing of CA1 Pyramidal Cells during Sharp Wave/Ripple Network Patterns in Exploratory Behavior. Neuron 49, 143–155 (2006).

25. T. K. Leonard, K. L. Hoffman, Sharp-Wave Ripples in Primates Are Enhanced near Remembered Visual Objects. Curr. Biol. 27, 257–262 (2017).

26. Y. Y. Chen, et al., Stability of ripple events during task engagement in human hippocampus. Cell Rep. 35, 109304 (2021).

27. Y. Norman, O. Raccah, S. Liu, J. Parvizi, R. Malach, Hippocampal ripples and their coordinated dialogue with the default mode network during recent and remote recollection. Neuron 109, 2767–2780.e5 (2021).

28. A. P. Vaz, S. K. Inati, N. Brunel, K. A. Zaghloul, Coupled ripple oscillations between the medial temporal lobe and neocortex retrieve human memory. Science 363, 975–978 (2019).

29. M. Nokia, J. Mikkonen, M. Penttonen, J. Wikgren, Disrupting neural activity related to awake-state sharp wave-ripple complexes prevents hippocampal learning. Front. Behav. Neurosci. 6, 84 (2012).

30. S. P. Jadhav, C. Kemere, P. W. German, L. M. Frank, Awake Hippocampal Sharp-Wave Ripples Support Spatial Memory. Science (2012) https://doi.org/10.1126/science.1217230 (January 12, 2022).

31. A. E. Papale, M. C. Zielinski, L. M. Frank, S. P. Jadhav, A. D. Redish, Interplay between Hippocampal Sharp-Wave-Ripple Events and Vicarious Trial and Error Behaviors in Decision Making. Neuron 92, 975–982 (2016).

32. C.-T. Wu, D. Haggerty, C. Kemere, D. Ji, Hippocampal awake replay in fear memory retrieval. Nat. Neurosci. 20, 571–580 (2017).

33. T. J. Davidson, F. Kloosterman, M. A. Wilson, Hippocampal replay of extended experience. Neuron 63, 497–507 (2009).

34. K. Diba, G. Buzsáki, Forward and reverse hippocampal place-cell sequences during ripples. Nat. Neurosci. 10, 1241–1242 (2007).

35. M. P. Karlsson, L. M. Frank, Awake replay of remote experiences in the hippocampus. Nat. Neurosci. 12, 913–918 (2009).

36. R. Todorova, M. Zugaro, Hippocampal ripples as a mode of communication with cortical and subcortical areas. Hippocampus 30, 39–49 (2020).

37. A. C. Singer, L. M. Frank, Rewarded Outcomes Enhance Reactivation of Experience in the Hippocampus. Neuron 64, 910–921 (2009).

38. A. T. Hussin, T. K. Leonard, K. L. Hoffman, Sharp-wave ripple features in macaques depend on behavioral state and cell-type specific firing. Hippocampus 30, 50–59 (2020).

39. A. M. Bastos, et al., Visual Areas Exert Feedforward and Feedback Influences through Distinct Frequency Channels. Neuron 85, 390–401 (2015).

40. D. Ferro, J. van Kempen, M. Boyd, S. Panzeri, A. Thiele, Directed information exchange between cortical layers in macaque V1 and V4 and its modulation by selective attention. Proc. Natl. Acad. Sci. 118 (2021).

41. G. G. Gregoriou, S. J. Gotts, H. Zhou, R. Desimone, High-Frequency, Long-Range Coupling Between Prefrontal and Visual Cortex During Attention. Science (2009) https://doi.org/10.1126/science.1171402 (January 17, 2022).

42. T. van Kerkoerle, et al., Alpha and gamma oscillations characterize feedback and feedforward processing in monkey visual cortex. Proc. Natl. Acad. Sci. 111, 14332–14341 (2014).

43. J. D. Semedo, A. Zandvakili, C. K. Machens, B. M. Yu, A. Kohn, Cortical Areas Interact through a Communication Subspace. Neuron 102, 249–259.e4 (2019).

44. J. Gold, J. Ciorciari, A neurocognitive model of flow states and the role of cerebellar internal models. Behav. Brain Res. 407, 113244 (2021).

45. H. Supèr, P. R. Roelfsema, Chronic multiunit recordings in behaving animals: Advantages and limitations. Prog. Brain Res. 147, 263–282 (2005).

46. , FMAToolbox (July 11, 2021).

47. E. Stark, et al., Pyramidal cell-interneuron interactions underlie hippocampal ripple oscillations. Neuron 83, 467–480 (2014).

48. K. H. Pettersen, A. Devor, I. Ulbert, A. M. Dale, G. T. Einevoll, Current-source density estimation based on inversion of electrostatic forward solution: Effects of finite extent of neuronal activity and conductivity discontinuities. J. Neurosci. Methods 154, 116–133 (2006).

49. N. K. Logothetis, C. Kayser, A. Oeltermann, In Vivo Measurement of Cortical Impedance Spectrum in Monkeys: Implications for Signal Propagation. Neuron 55, 809–823 (2007).

50. V. B. Mountcastle, Modality and topographic properties of single neurons of cat’s somatic sensory cortex. J. Neurophysiol. 20, 408–434 (1957).

51. C. E. Schroeder, C. E. Tenke, S. J. Givre, J. C. Arezzo, H. G. Vaughan, Striate cortical contribution to the surface-recorded pattern-reversal vep in the alert monkey. Vision Res. 31, 1143–1157 (1991).

52. S. J. Givre, C. E. Schroeder, J. C. Arezzo, Contribution of extrastriate area V4 to the surface-recorded flash VEP in the awake macaque. Vision Res. 34, 415–428 (1994).

53. M. A. Gieselmann, A. Thiele, Stimulus dependence of directed information exchange between cortical layers in macaque V1. bioRxiv, 2020.07.10.197566-2020.07.10.197566 (2020).

54. H. Bokil, P. Andrews, J. E. Kulkarni, S. Mehta, P. P. Mitra, Chronux: A platform for analyzing neural signals. J. Neurosci. Methods 192, 146–151 (2010).

55. M. X. Cohen, A better way to define and describe Morlet wavelets for time-frequency analysis. NeuroImage 199, 81–86 (2019).

56. G. Buzsáki, et al., Hippocampal network patterns of activity in the mouse. Neuroscience 116, 201–211 (2003).

57. Y. Benjamini, Y. Hochberg, Controlling the False Discovery Rate: A Practical and Powerful Approach to Multiple Testing. J. R. Stat. Soc. Ser. B Methodol. 57, 289–300 (1995).

58. A. Thiele, L. S. Delicato, M. J. Roberts, M. A. Gieselmann, A novel electrode-pipette design for simultaneous recording of extracellular spikes and iontophoretic drug application in awake behaving monkeys. J. Neurosci. Methods 158, 207–211 (2006).

59. H. Gray, et al., Physiological, behavioral, and scientific impact of different fluid control protocols in the rhesus macaque (Macaca mulatta). eNeuro 3 (2016).

60. M. A. Gieselmann, A. Thiele, Comparison of spatial integration and surround suppression characteristics in spiking activity and the local field potential in macaque V1. Eur. J. Neurosci. 28, 447–459 (2008).

61. P. R. Roelfsema, M. Tolboom, P. S. Khayat, Different Processing Phases for Features, Figures, and Selective Attention in the Primary Visual Cortex. Neuron 56, 785–792 (2007).

